# The glucocorticoid receptor elicited proliferative response in human erythropoiesis is BCL11A-dependent

**DOI:** 10.1101/2024.02.05.577972

**Authors:** Maria Mazzarini, Jennifer Cherone, Truong Nguyen, Fabrizio Martelli, Lilian Varricchio, Alister P.W. Funnell, Thalia Papayannopoulou, Anna Rita Migliaccio

## Abstract

Prior evidence indicates that the erythroid cellular response to glucocorticoids (GC) has developmental specificity, namely, that developmentally more advanced cells that are undergoing or have undergone fetal to adult globin switching are more responsive to GC-induced expansion. To investigate the molecular underpinnings of this, we focused on the major developmental globin regulator BCL11A. We compared: **a)** levels of expression and nuclear content of BCL11A in adult erythroid cells upon GC stimulation; **b)** response to GC of CD34+ cells from patients with *BCL11A* microdeletions and reduced *BCL11A* expression, and; **c)** response to GC of two cellular models (HUDEP-2 and adult CD34+ cells) before and after reduction of *BCL11A* expression by shRNA. We observed that: **a)** GC-expanded erythroid cells from a large cohort of blood donors displayed amplified expression and nuclear accumulation of BCL11A; **b)** CD34+ cells from *BCL11A* microdeletion patients generated fewer erythroid cells when cultured with GC compared to their parents, while the erythroid expansion of the patients was similar to that of their parents in cultures without GC, and; **c)** adult CD34+ cells and HUDEP-2 cells with shRNA-depleted expression of *BCL11A* exhibit reduced expansion in response to GC. In addition, RNA-seq profiling of shRNA-BCL11A CD34+ cells cultured with and without GC was similar (very few differentially expressed genes), while GC-specific responses (differential expression of *GILZ* and of numerous additional genes) were observed only in controls cells with unperturbed BCL11A expression. These data indicate that BCL11A is an important participant of certain aspects of the stress pathway sustained by GC.

## INTRODUCTION

For many decades, corticosteroids have been used for the treatment of patients with certain anemias [1,2] and in vitro studies have reported the preferential effects of steroids in modulating human erythroid cells [3]. More recently, the effect of corticosteroids on erythropoiesis has been extensively explored using murine model systems[4]. It has emerged that glucocorticoids (GC) are required for expansion of erythroid progenitor cells, especially under stress conditions [5,6]. Under these conditions, GC also cooperate with cytokine membrane receptors, such as stem cell factor (SCF) and erythropoietin (EPO) receptors, that are also essential for stress responses [7,8], and with other participating signaling molecules (Zinc Finger Protein 36, C3H1 Type-Like 2, Peroxisome proliferator-activated receptor α and p57Kip2) [9–11]. Later studies indicated that in humans, GC additionally act on more mature cells that extend beyond the BFU-E stage[12], including CFU-E and pro-erythroblasts which, in contrast to cells generated without GC, retain expression of cKIT, the receptor for SCF [11]. Additional markers for human stress erythropoiesis (erythroferrone, growth differentiation factor 15, hemoglobin subunit gamma 2, HBG2) have also been identified [13,14]. The distinct profiles of human erythroid progenitors generated in the presence of GC were also uncovered [15] and it was found that, in addition to CD34, GC-generated progenitor cells express MPL, the receptor for thrombopoietin, and CD36, the thrombospondin receptor [16,17].

Despite these advances in understanding on the target cells involved (mainly murine BFU-E and human BFU-E/CFU-E) and the several participating pathways activated under stress conditions, the mechanistic molecular details of GC effects on erythropoiesis have remained poorly delineated. Furthermore, an apparent developmental specificity in response to GC, also remains unexplored. Specifically, the proliferative response to GC differs markedly between two widely used cord blood-derived erythroid cell lines: HUDEP-1 and HUDEP-2[18]. HUDEP-1 cells, which express fetal globins (*HBG1/2*), do not respond to GC, in contrast to their counterparts, HUDEP-2, which have undergone globin switching and express the adult forms (*HBB1/2)*. Relatedly, cells originating from some cord blood units have elsewhere been reported to be unresponsive to GC [11], in contrast to the consistent effect that is observed in cells from adult sources [16]. Moreover, hybrid cells generated by fusing human fetal liver-derived and adult murine MEL cells express human fetal globin for several weeks in culture, but eventually switch to an adult phenotype [19]. Consistent with the above, a GC response was only observed in these cells during and following the switching period, and not prior to this, and interestingly, this response was accompanied by an acceleration of the *HBG* to *HBB* switch [20]. These intriguing prior data inspired us here to explore whether the key erythroid and globin switching regulator BCL11A might play a role in the developmental stage specificity of the GC response in human erythropoiesis.

## MATERIALS AND METHODS (see Supplementary and Table S1 for detail)

### Human subjects

Blood from 25 normal donors and from three heterozygous *BCL11A-* microdeletion patients [21] and their parents was provided by the Italian Red Cross Blood Bank (Rome, Italy) and Perugia and Palermo University (Italy) respectively. Human CD34+ cells were purchased from the Fred Hutchinson Cancer Center (Seattle, WA, USA).

### Proliferation (Prol) and differentiation (Diff) erythroid cultures

Blood mononuclear cells (10^6^cells/mL) cultured for 10 days with SCF (10ng/mL), EPO (1U/mL), interleukin-3 (IL-3, 1ng/mL), dexamethasone (Dex, 10^−6^M) and estradiol (10^−6^M) (Prol) and their progeny were induced to differentiate (Diff) for 4 days with EPO (3U/mL) and insulin (10ng/mL) [22].

### Colony assays

Blood mononuclear cells (3×10^5^cells/mL) CD34+ cells (6×10^3^cells/mL) and their progeny were cultured, respectively, in MethoCult and MethoCult™ H4434. Colonies derived from BFU-E, CFU-GM, and CFU-GEMM were identified according to standard morphological criteria [23].

### RNA isolation and quantitative real-time PCR

Total RNA was isolated from 10^6^ cells using Trizol and reverse transcribed with RNase OUT kits [24]. Quantitative real-time PCR was performed as described in the Supplementary (**Figure S1**). To assess *BCL11A* knockdown, RNA was extracted with the Qiagen RNeasy Micro Kit and reverse transcribed with the iScript Reverse Transcription Supermix. mRNA levels and expression ratios were calculated as described in the Supplementary.

### Western Blot analysis

Whole cell extracts (30μg protein/lane) and nuclear protein fractions (15μg/lane) were separated on SDS-PAGE and transferred to nitrocellulose membranes which were probed with antibodies against BCL11A, GATA1, HBB, HBG, GRα, GAPDH, Lamin B1 and Histone 3 and then with horseradish peroxidase-coupled secondary antibodies.

### Flow cytometry

Erythroid differentiation was assessed by flow cytometry with Phycoerythrin (PE)-CD34, Pacific Blue 450 (PB450)-CD36 and fluorescein isothiocyanate (FITC)-CD235a. Data were analyzed with FlowJo.

### Knockdown of *BCL11A* in HUDEP-1 and HUDEP-2 cells

Cell lines were cultured as described [18] and transduced (1×10^5^cells/0.4μL of virus) either with the shRNA *Luciferase* (*Luc*)(*shLuc*, control) or the commercial *BCL11A* shRNA (*shBCL11A*).

### Knockdown of *BCL11A* in CD34+ cells

CD34+ cells were cultured for 2 days in expansion media (StemSpan H3000) and transduced on Day -5 with either *shLuc* or *shBCL11A* (1×10^5^cells/3.2μL of virus)(**Figure S2),** as described in the Supplementary [25]. Transduced cells were selected for 3 days with puromycin (0.5μg/mL) and 2×10^5^cells/mL were transferred at Day 0 to erythroid expansion media plus/minus hydrocortisone (1μM). Preliminary experiments indicated that Hydrocortisone is equally potent to Dex in expanding CD34+ cells (**Figure S3**).

### RNA-sequencing (RNAseq)

RNAseq of BCL11A-microdeletion patients was performed as described[21]. The day 3 and day 5 progeny of CD34+ cells were profiled with the Illumina libraries constructed using Prime-seq [26] and sequenced at read lengths 28×8×94. RNA-seq mapping, demultiplexing and quantification were performed with STARsolo using GENCODEv25 for annotation, featureCounts for gene counts and DESeq2 to identify differentially expressed genes (DEGs) between groups.

### GeneSet Enrichment Analysis

GSEA was performed with GSEA 4.3.1 with default parameters. Expression pathways were analyzed with the Hallmark, KEGG and REACTOME gene-set.

### Statistical Analyses

Results are presented as median, 25%-75% interquartile range and maximum/minimum or as Mean(+/-SEM), as appropriate. Results were compared by Wilcoxon signed-rank test, paired t-test or Anova multiple test, as appropriate, using GraphPad Prism 9.

### Data Sharing Statement

The RNAseq data will be deposited in publicly available databases upon the acceptance of the manuscript.

## RESULTS

### GC receptor activation increases expression and nuclear content of BCL11A in Erys expanded *in vitro* from healthy CD34+ cells

As reported, blood mononuclear cells from normal donors generate a population of macrocytic erythrocytes when cultured for 10 days with Dex (“Prol”, **Figure 1A**) and then acquire a mature morphology, including a size reduction, after exposure for 4 days to EPO (“Diff”, **Figure 1A**) [22]. This maturation is associated with significant increases (∼50%) in total mRNA content, in large part due to a >10-fold increase in adult *HBA* and *HBB* globins, as well as a 5-fold increase in *AHSP* transcripts [24]. There is a concomitant decrease in the average fetal-to-adult globin expression ratio (*HBG/(HBG+HBB))* from 0.4±0.2 at the day 10 Prol stage to 0.08±0.01 (p<0.05) in Diff cultures [24].

**Figure 1.**
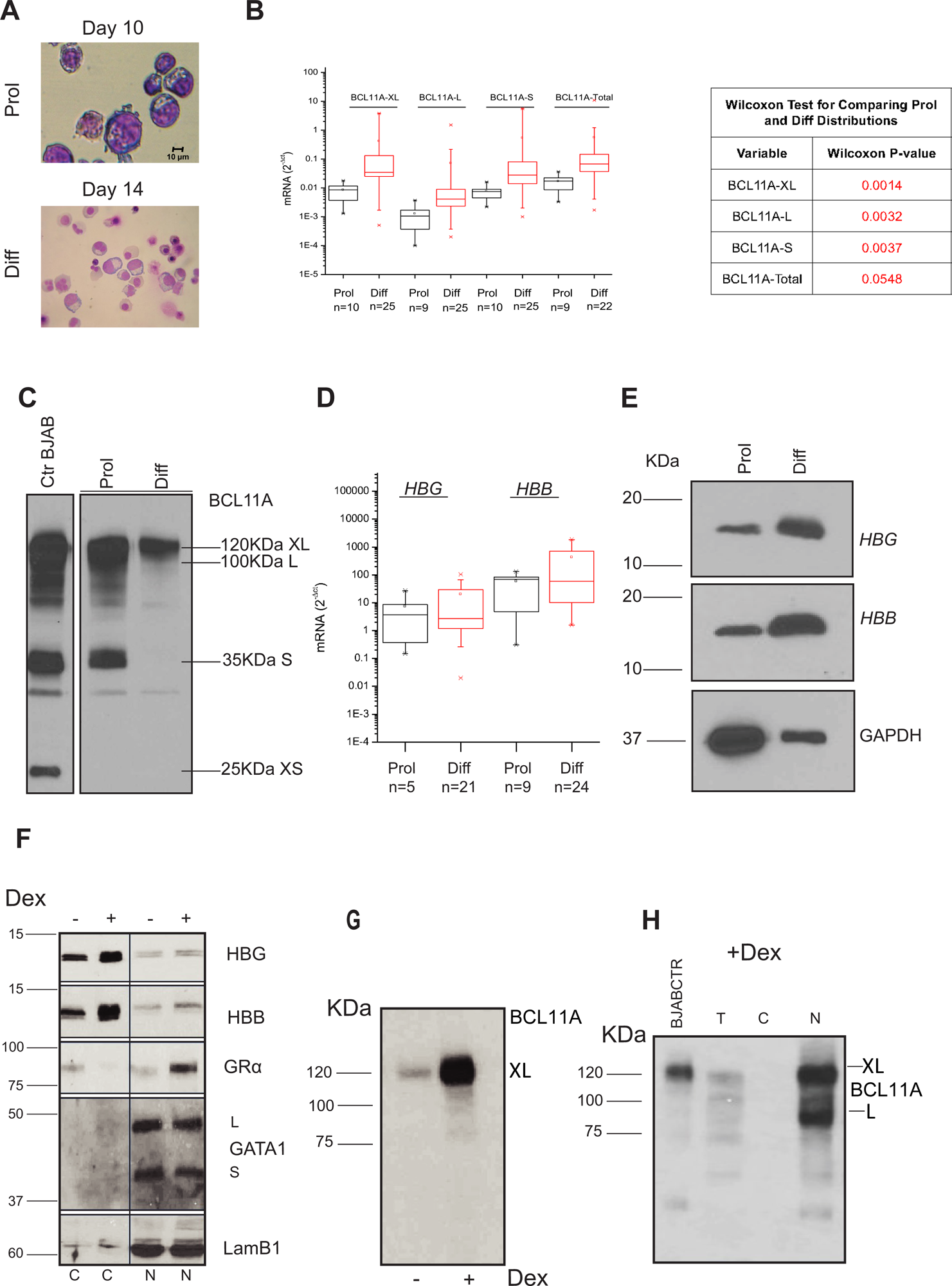
Erythroid cells generated ex vivo in the presence of Dex from a large cohort (n=25) of adult healthy donors (AB) express high levels of *BCL11A* and *globin* genes. **A)** Representative morphology by May-Grunwald staining of Erys obtained at day 10 of cultures with Dex (Prol) and subsequently induced to mature (Diff) for 4 days with EPO. **B)** Quantitative RT-PCR determinations of the levels of mRNA for the total and *BCL11A* isoforms expressed by Erys at day 10 of Prol culture and by Erys after 4 days of Diff culture. Values are expressed as median, 25% to 75% interquartile range and maximum and minimum of values. The number of individual subjects included in each group is indicated by n. p-values among the levels of the *BCL11A* mRNA isoforms expressed in Prol and Diff are presented in the Table on the right. **C)** Western Blot determinations of the BCL11A content of Erys from AB obtained in Prol and Diff cultures. The lysate extract from BJABl cells derived from Burkit lymphoma[35] was used as a positive control. **D)** Levels of *HBG* and *HBB* expressed by the cells in Prol and Diff (the same donors presented in A). These results were already published[24] and are presented only for comparison. Differences among groups are not statistically significant. **E)** Representative HBG and HBB protein content in Erys under Prol and Diff conditions. GAPDH was used as loading control. **F)** Western Blot analysis of GRα content in extracts containing the cytoplasmic (C) and nuclear (N) proteins of Erys obtained by day 10 in cultures with and without Dex as indicated. The content of globins and GATA1 was analyzed for comparison. Lamin B1 was analyzed as loading control of the nuclear fraction. The position of the molecular weights (kDa) is indicated on the left. **G)** Content of BCL11A in Erys obtained at day 10 in cultures without and with dexamethasone (Dex). **H)** Western Blot analysis of BCL11A content in the total (T), cytoplasmic (C) and nuclear fractions (N) from Erys expanded with Dex. BJAB total extract was loaded as a positive control. The fact that BCL11A was more readily detectable in nuclear extracts than in total cell extracts reflects the relative concentration of the protein in the two samples.

Erys generated from different donors were found to express widely different levels of the differentially spliced *BCL11A* mRNA isoforms extra-long (XL), long (L) and small (S) (**Figure 1B**), spanning almost 4 orders of magnitude. All three isoforms showed a consistent increase in expression across donors upon differentiation (**Figure 1B**). The *BCL11A* isoforms that increase most significantly in expression between immature and mature Erys are the full-length XL and L isoforms, consistent with previous reports [27]. At the protein level, all three BCL11A isoforms are expressed by the immature population (**Figure 1C**). Following differentiation, cells express mainly BCL11A-XL (**Figure 1C**). This isoform shift is associated with increases in *HBG* and *HBB* expression at the mRNA (**Figure 1D**) and protein (**Figure 1E**) levels.

The connection between BCL11A expression and GC receptor (GR) activation was explored by determining the effects of Dex on the total and nuclear (as a surrogate of nuclear activity) content of BCL11A in Erys expanded in culture from healthy donors. We confirmed that Dex leads to GR activation under these conditions by determining that, adult Erys generated with Dex contain greater levels of GR in the nucleus than those generated without Dex, while the nuclear content of GATA1 remains similar (**Figure 1F**). By contrast, the cytoplasmic content of GR is decreased by Dex, while that of the globin chains is increased. Interestingly, the total cellular content of BCL11A, and specifically of the XL isoform, is much greater in cells generated with Dex than in those without (**Figure 1G)** with robust localization in the nuclear fraction (**Figure 1H**). These data suggest that in Erys cultured with Dex, BCL11A is mostly in the nucleus and, therefore, presumably active.

### Erythroid progenitors from *BCL11A*-deficient patients respond poorly to Dex

To further assess the relationship between GR and *BCL11A* in stress erythropoiesis, we analyzed the response to Dex *in vitro* of hematopoietic progenitor cells from the blood of three patients carrying microdeletions spanning the *BCL11A* locus [21]. Cells from their parents (Mother + Father) were used as controls.

The CFC (**Figure 2A**) and the frequency of CD34+ cells (0.6±0.3 vs 0.28±0.2%, respectively) present in the blood from the patients had a lower trend but were not statistically different from that of their parents. By day 10, after Dex treatment, erythroid progenitor cells from the patients generated significantly lower numbers of Erys than those from their parents (**Figure 2B**). In addition, while as expected, Erys generated by the parents with Dex are mostly immature, those generated by the patients have a more mature morphology (**Figure 2C**). In the case of patient 2, we determined that the number of Erys generated by the patient is similar to that generated by the mother in the absence of Dex but is lower than that generated by the mother in cultures with Dex (**Figure 2D**).

**Figure 2.**
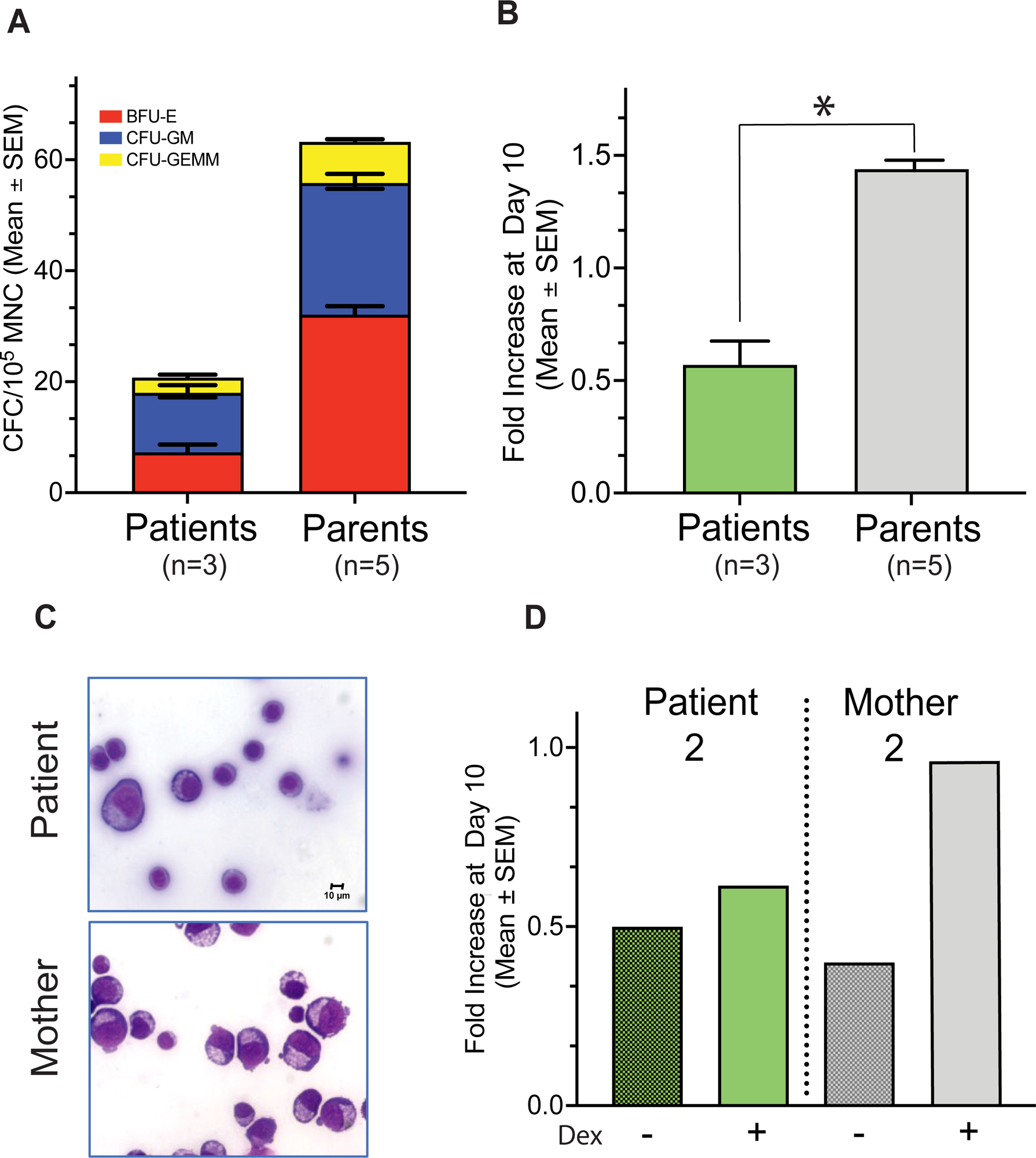
The blood from BCL11A-deficient patients contain normal numbers of hematopoietic progenitor cells but these cells generate low numbers of Erys when cultured with Dex. **A)** Number of colonies from 10^5^ blood mononuclear cells (MNC) from the patients and the parents. **B)** Fold increase (FI) with respect to day 0, observed at day 10 in cultures of BCL11A-deficient patients and of their parents stimulated with Dex. **C)** Representative morphology by May-Grunwald staining of Erys obtained at day 10 of cultures with Dex from patient 2 and from the mother. **D)** Fold increase (FI) with respect to day 0, generated at day 10 by patient 2 and his mother in parallel cultures stimulated with and without Dex. In A, B and D, results are presented as Mean (±SEM) of values observed in multiple individuals per each group and statistically significant p values (p<0.05) among groups calculated by Anova are indicated by asterisk. The number of individuals included in each group is indicated by n.

Lastly, RNAseq data confirmed that Erys generated from the three patients express lower levels of *BCL11A* (associated with higher fetal-to-adult *HBG:HBB* levels) than Erys generated by the parents (**Figure 3**). Specifically, the levels of *BCL11A* expressed by Erys from patient 2, which responded poorly to Dex (**Figure 2D**), were barely detectable (**Figure 3**).

**Figure 3.**
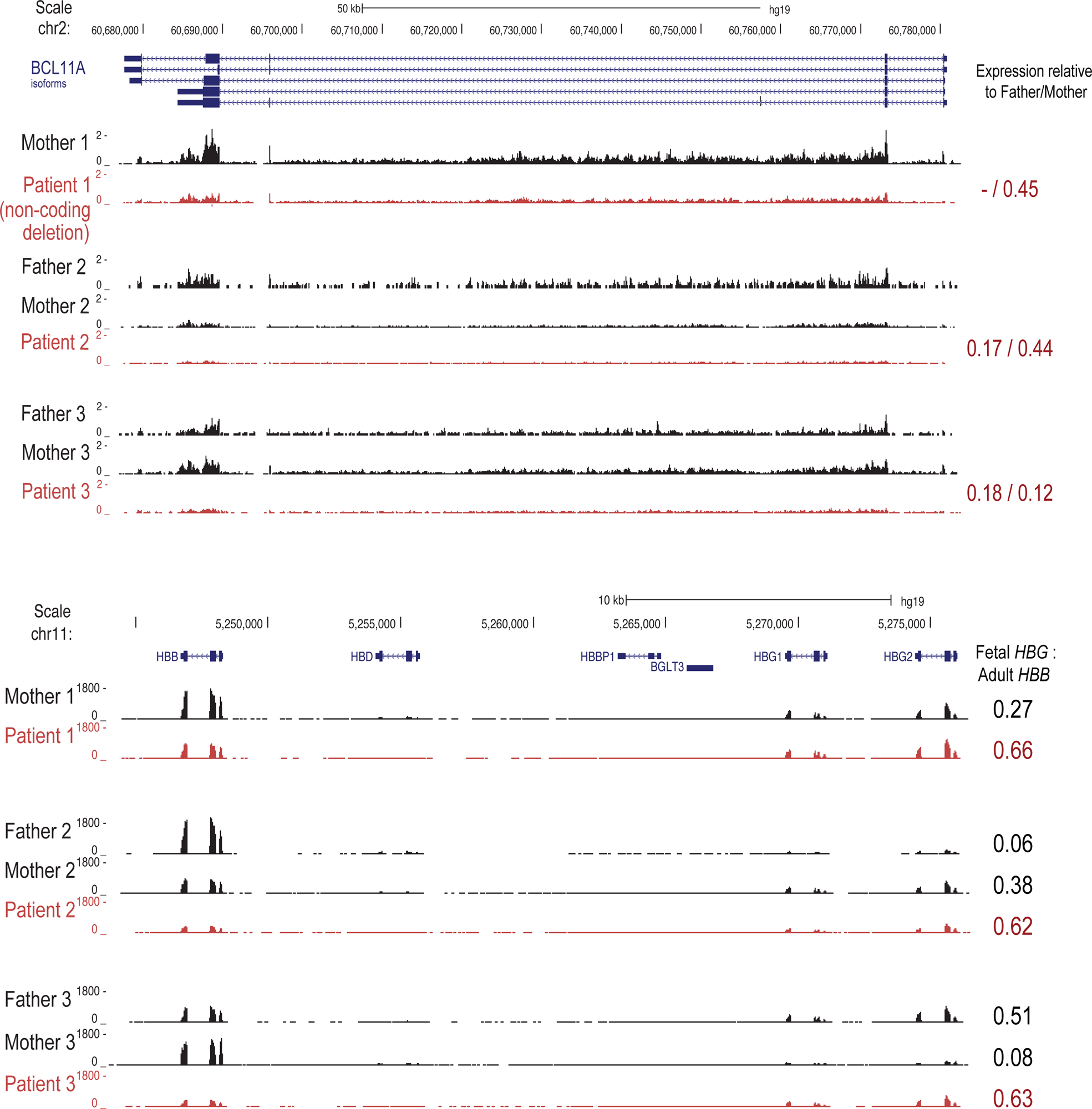
Erythroid cells expanded from BCL11A-deficient patients express reduced levels of *BCL11A* and increased levels of *HBG* mRNA. RNAseq analyses for *BCL11A*, *HBB* and *HBG* mRNA expression in Erys generated at day 10 by patient 1, carrying a 3.5 Mb deletion downstream to *BCL11A* encompassing the erythroid specific enhancer, and patients 2 and 3, carrying a 642 kb and 2.5-Mb deletions on the BCL11A gene, respectively. Results obtained by sequencing the mRNA from Erys generated in parallel cultures by the parents of the patients are presented for comparison.

### Loss of function of *BCL11A* reduces the response to GC of HUDEP-2 and CD34+ cells

As previously reported[18], immortalized Erys with a fetal phenotype (namely, HUDEP-1 cells expressing fetal globin) do not respond to Dex, in contrast to their adult phenotype counterpart (HUDEP-2) which do respond (**Figure 4A**). The *BCL11A* shRNA reduced to barely detectable levels of the *BCL11A* mRNA and nuclear protein content in HUDEP-2 cells (**Figure 4B-C**). Consequentially, *HBG* levels increased by 2-fold with respect to those observed in the shRNA control and non-treated HUDEP-2 cells (**Figure 4B**). Interestingly, the HUDEP-2 cells with low *BCL11A* expression were found to not respond to GC treatment (**Figure 4A**).

**Figure 4.**
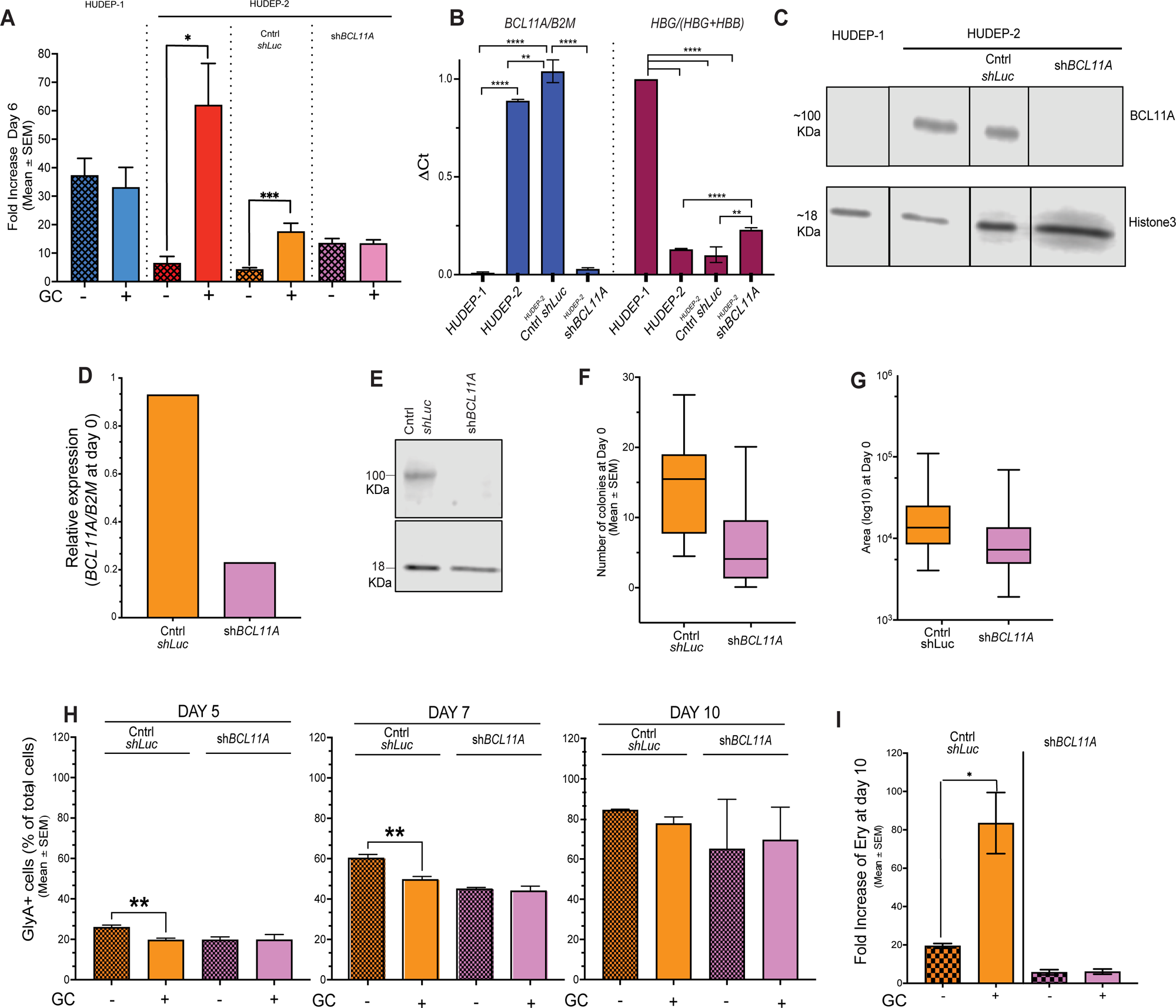
GC do not improve the number of erythroid cells generated in culture by progenitor cells in which *BCL11A* has been knocked down by a shRNA approach. **A)** Fold increase in the number of cells observed after six days of culture of untreated HUDEP-1 and HUDEP-2 cells and of HUDEP-2 cells transduced with either the control or the *shBCL11A* vector with and without GC. **B)** Quantitative RT-PCR for the expression of *BCL11A* and *HBG* in untreated HUDEP-1 and HUDEP-2 cells and in HUDEP-2 cells transduced with either the control or the *shBCL11A* vector, as indicated. **C)** Western Blot analysis of the BCL11A content in nuclear extracts from untreated HUDEP-1 and HUDEP-2 cells and from HUDEP-2 cells transduced with either the control or the *shBCL11A* vector, as indicated. **D)** Quantitative RT-PCR and **E)** Western Blot analyses for the expression of *BCL11A* by CD34+ cells transduced with either the control *shLuc* or the sh*BCL11A* vector. **F)** Number of BFU-E and **G)** size of the colonies generated by CD34+ cells transduced with the control *shLuc* or *shBCL11A* vector. Results are presented as median, 25% to 75% interquartile range and maximum and minimum range. The cells analyzed in A-D are at Day 0 of the erythroid expansion culture. **H)** Frequency of GlyA+ cells generated by either the control *shLuc* or the sh*BCL11A* CD34+ cells at day 5, 7 and 10 of erythroid cultures with and without GC. Representative flow cytometry dot plots are presented in **Figure S4**. Fold increase in the number of cells generated at day 10 of culture by control *shLuc* and sh*BCL11A* CD34+ cells with or without GC. Results are presented as Mean (±SEM) of values observed in three independent experiments. p values among groups are indicated by asterisks within the panels (*, p<0.05;**,p<0.001; ****; p<0.0001).

To investigate the effect of decreased *BCL11A* expression in normal adult Erys, we transduced CD34+ cells with either the sh*BCL11A* or the control sh*Luc* vector. The sh*BCL11A* CD34+ cells express reduced levels (∼70%) of *BCL11A* mRNA **(Figure 4D)** and barely detectable nuclear protein with respect to controls **(Figure 4E)**.

The numbers and sizes of erythroid colonies from CD34+ with *BCL11A* shRNA showed a lower trend which did not reach statistical significance compared to control **(Figure4 F, G).** As expected, we observed that control cells, upon GC stimulation, generated reduced numbers of GlyA (CD235a)^pos^ cells at early days (5 and 7) of culture (**Figures 4H,S4)** which was associated with greater numbers of total Erys observed by day 10 (**Figure 4I)**, an indication that GC delayed differentiation favoring expansion of control cells. In contrast, the sh*BCL11A* CD34+ cells generated similar numbers of GlyA positive cells and of total cells with and without GC (**Figure4 H,I).** The reduced Ery expansion from sh*BCL11A* vs control cells without GC is not statistically significant.

These results indicate that loss or significant reduction of *BCL11A* expression reduces the response of erythroid progenitors to GC.

### BCL11A downregulation impairs the molecular response of erythroid cells to GC

To investigate the molecular consequences of GC stimulation on erythroid cells, we performed RNAseq on samples taken at days 3 and 5 of culture with and without GC. Day 3 is when GC induces CD34+ cells to generate stress-specific BFU-E [17] while day 5 is when CD34+ cells are no longer detectable **(Figure S4)** and CFU-E prevail [11]. Hierarchical clustering of Euclidean distances and PCA analysis indicated clear segregation between control and *BCL11A* shRNA cells both at day 3 and 5 (**Figure S5A,B**). There is also clear segregation between cells (control and sh*BCL11A*) treated with and without GC at day 3 while the difference between these two groups at day 5 is less clear (**Figure S5B**). In the control cells, the statistically significant (padjust <0.05) DEGs between plus or minus GC cells were 686 at day 3 (259 upregulated, 427 downregulated) and 245 (78 upregulated, 167 downregulated) at day 5 (**Figure S5C**). By contrast, there were only 191 DEGs (91 upregulated, 100 downregulated) in sh*BCL11A* cells plus and minus GC at day 3 and only 20 DEGs (17 upregulated, 3 downregulated) at day 5 (**Figure S5C**). Of the 20 DEGs observed at day 5 in the sh*BCL11A* group, 11 are also significantly affected by Dex in the control group (**Table S2**). Of interest, one of the genes downregulated by GC both in control and sh*BCL11A* cells at day 5 is cKIT, a possible reflection of the increased cKIT metabolism following SCF-cKIT interactions in erythroid precursors generated in the presence of GC [11].

Furthermore, the RNAseq data indicated that the ubiquitous GR target *GILZ* was maximally activated at day 3 with a significant difference between the control and shRNA group (**Figure 5A**). Further comparison of the expression of GR targets genes available in a public human data base (https://classic.wikipathways.org/index.php/Pathway:WP2880#nogo2) indicated that GR altered the expression of 10 (*CAVIN2, ALOX5AP, S100P, GADD45B, SRGN, SERPINB9, RGS2, SPRY1, AMIGO2, CCL2* at day 3) and 1 (*CCL2*, at day 5) genes in the control group vs only 3 (*RGS2, SPRY1, CCL2* at day 3) and none (at day 5) in the shRNA group (**Figure 5B**). These genes are involved in inflammatory, growth and apoptosis responses to GC.

**Figure 5.**
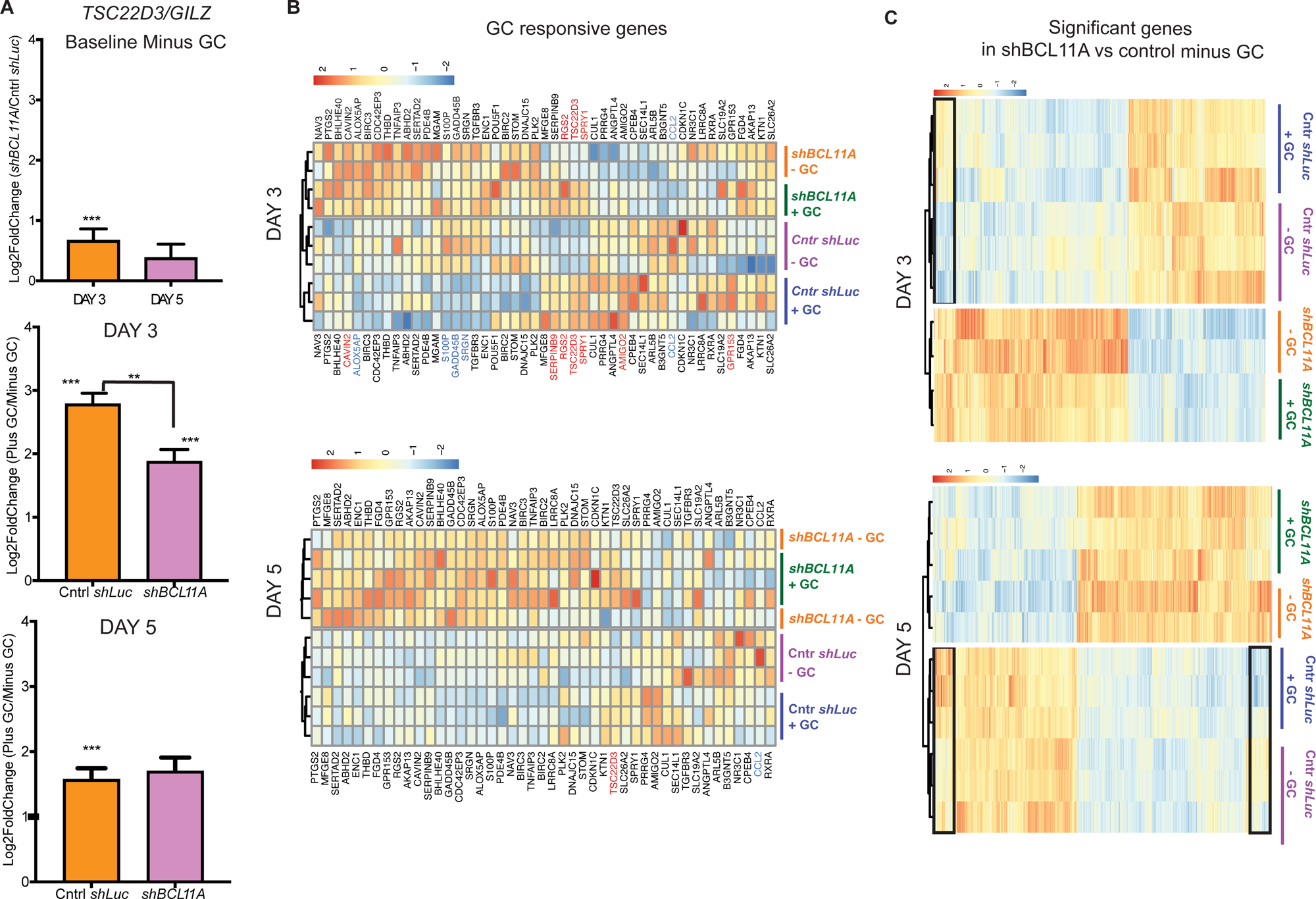
GC activates the expression of a limited number of GC target genes in progeny generated by *shBCL11A* CD34+ cells at day 3 and day 5 of erythroid expansion. **A)** log2Fold change (± lfcSE) in the expression of the ubiquitous GC-target gene *GILZ (TS22D3)*. Top panel: comparison of the baseline expression of *GILZ* in the progeny of sh*BCL11A* and control *shLuc* CD34+ cells cultured without GC. Middle and bottom panel: comparison of the expression of *GILZ* in the progeny of sh*BCL11A* and control *shLuc* CD34+ cells cultured with and without GC for 3 and 5 days as indicated. **B)** Heatmaps of GC responsive genes described in a publicly available data base (https://classic.wikipathways.org/index.php/Pathway:WP2880#nogo2) in control *shLuc* and sh*BCL11A* CD34+ cells cultured with or without GC for 3 and 5 days. Red and blue fonts indicate statistically significant up- and down-regulated genes, respectively. **C)** Heatmaps of all the genes expressed at significantly different levels in control *shlLuc* and sh*BCL11A* CD34+ cells cultured without GC for 3 and 5 days. The black boxes indicate genes that are expressed at significantly different levels in *shLuc* cultured with and without GC. See **Figures S5** and **Tables S2-S5** for further detail.

Pathway analyses of all the genes differentially expressed among groups (**Figure 5C)** identified several signaling activated by GC in control cells (**Figure 6A**). As an example, the KEGG cured gene set identified downregulation of the hematopoietic cell signature, both at day 3 (NES=-1.91, fdrq=0.000) and day 5 (NES=-1.94, fdrq=0.001), and upregulation of genes involved in protein biosynthesis by day 5 only, supporting the hypothesis that GR retained control erythroid cells immature and upregulates protein translation. In addition to pathways involved in protein biosynthesis, reactome analyses identified that GC also activates genes associated with mitochondrial activity in control cells (**Figure S6**). GSEA analysis identified in these GC-treated control cells a strong downregulation of pathways involved in inflammatory and immune responses (**Figures 6A**, **S6, S7**). By contrast, the GO enrichment program and pathways analyses identified that in sh*BCL11A* cells, GC mostly affected pathways related to inflammation and immune responses (**Figure 6B**).

**Figure 6.**
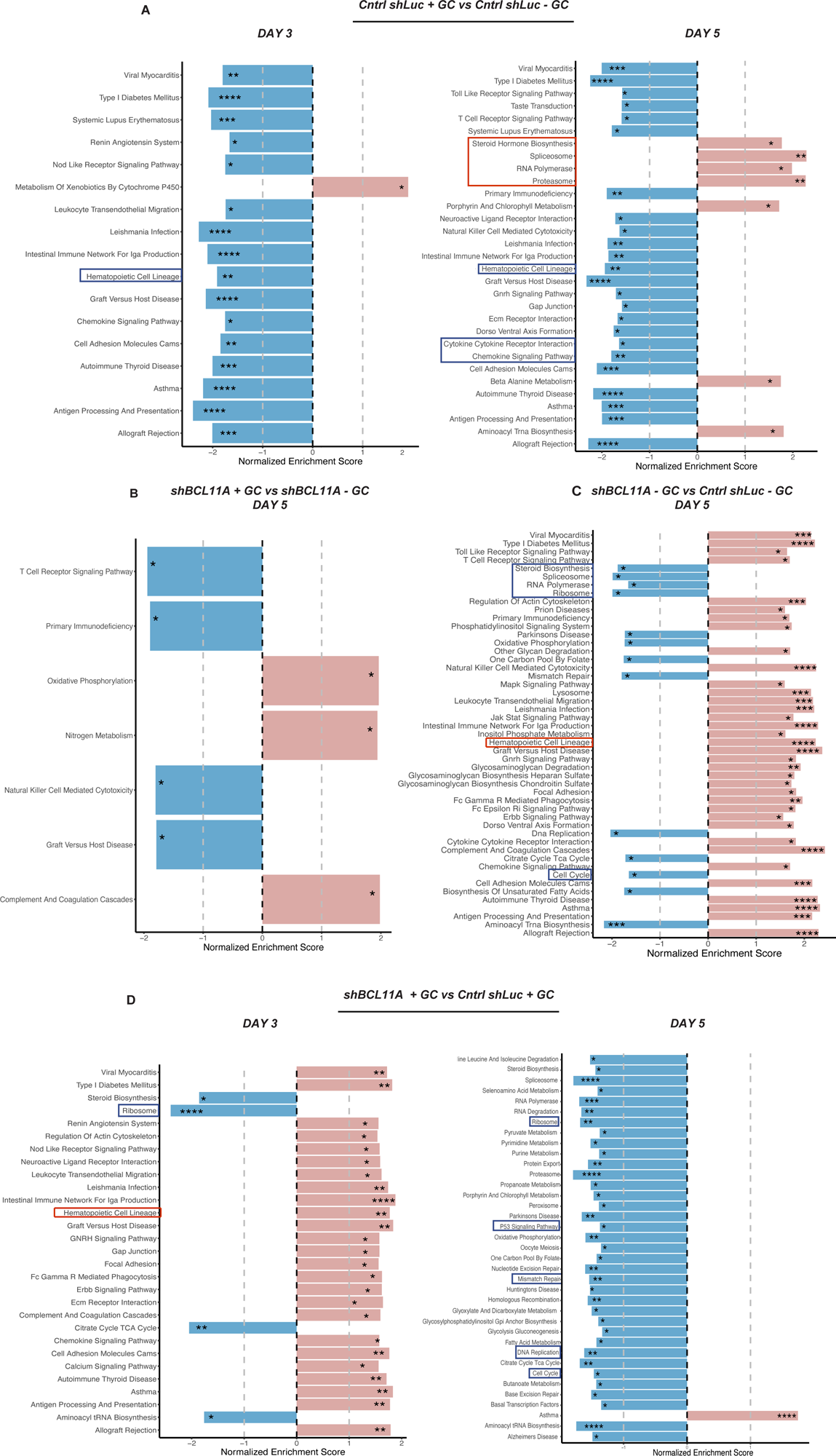
Glucorticoid stimulation regulates pathways of the hematopoietic cell lineage and in protein biosynthesis and this activity is blunt in the absence of BCL11A. GSEA comparison between control *shLuc* cells with or without GC at day 3 and day 5 **A)**, and between *shBCL11A* with or without GC (**B**), between *shBCL11A* and *shLuc* without GC (**C**) at day 5 and with GC addition (**D**) at day 3 and day 5. Red and blue columns indicate enriched and un-enriched pathway, respectively. All the data were compared using the KEGG cured gene set (c2.cp.kegg.v2023.1.Hs.symbols.gmt). Red and blue boxes indicate respectively up- and down-enriched pathways. FDR q-values among groups are indicated by asterisks (*, p<0.05; **,p<0.001; ***; p<0.0001****; p<0.00001).

Overall, the greatest variability in gene expression was observed between control and sh*BCL11A* cells cultured with and without GC (**Figure S5C**). Specifically, control and sh*BCL11A*-cells cultured without GC differentially expressed, in addition to *HBG* and *BCL11A* (**Table 1**), 1719 genes (1090 upregulated, 629 downregulated) at day 3 and 1939 (1273 upregulated, 666 downregulated) at day 5 (**Figure S5C**). In addition, the number of DEGs statistically significant between sh*BCL11A* and shLuc cells plus GC were 2,077 at day 3 (1,989 upregulated, 818 downregulated) and 2,100 at day 5 (1281 upregulated, 819 downregulated) (**Figure S5C**).

**Table 1:** Selected genes differentially expressed in control *shLuc* cells with and without GC and in sh*BCL11A* cells vs control *shLuc* cells with and without GC. Genes were considered upregulated and downregulates with a log2Fold Change > 0.500 (red fonts) or < - 0.500, respectively. Bold font indicates the statistically significant genes (padj < 0.05).

|  | Gene ID | Cntl *shLuc*:<br>Plus GC vs Minus GC |  |  |  | Minus GC:<br>*shBCL11A* vs cntl *shLuc* |  |  |  | Plus GC:<br>*shBCL11A* vs cntl *shLuc* |  |  |  |
| --- | --- | --- | --- | --- | --- | --- | --- | --- | --- | --- | --- | --- | --- |
|  |  | DAY 3 |  | DAY 5 |  | DAY 3 |  | DAY 5 |  | DAY 3 |  | DAY 5 |  |
|  |  | log2<br>Fold<br>change | padj | log2<br>Fold<br>change | padj | log2<br>Fold<br>change | padj | log2<br>Fold<br>change | padj | log2<br>Fold<br>change | padj | log2<br>Fold<br>change | padj |
| GLOBINS | *HBG2* | 0.055 | 0.983 | 0.094 | 0.623 | -0.153 | 0.522 | -0.52 | 0 | -0.305 | 0.038 | -0.685 | 0.000 |
|  | *HBG1* | -0.087 | 0.935 | 0.041 | 0.869 | -0.066 | 0.821 | -0.371 | 0.001 | -0.122 | 0.501 | -0.520 | 0.000 |
|  | *HBB* | -0.035 | 0.986 | -0.098 | 0.639 | -0.486 | 0.001 | -0.823 | 0 | -0.658 | 0.000 | -0.879 | 0.000 |
| TRANSCRIPTION<br>FACTORS | *KLF1* | -0.05 | 0.986 | 0.125 | 0.464 | -0.363 | 0.084 | -0.114 | 0.472 | -0.342 | 0.062 | -0.624 | 0.000 |
|  | *BCL11A* | -0.03 | 0.993 | -0.185 | 0.472 | -1.19 | 0 | -1.291 | 0 | -0.909 | 0.000 | -1.186 | 0.000 |
| CELL CYCLE<br>INHIBITORS | *CDKN2D* | -0.494 | 0.008 | -0.386 | 0.205 | 1.154 | 0 | 0.853 | 0 | 1.137 | 0.000 | 1.179 | 0.000 |
|  | *CDKN2B* | 0.483 | 0.572 | 0.795 | NA | 0.763 | 0.069 | 1.601 | 0.129 | 0.956 | 0.012 | 2.389 | 0.000 |
| QUIESCENCE<br>REGULATORS | *EGR1* | -0.142 | 0.888 | -0.499 | 0.193 | -0.203 | 0.445 | 0.16 | 0.747 | 0.323 | 0.306 | 1.523 | 0.000 |
|  | *FOS* | -0.42 | 0.129 | -0.681 | 0.002 | -0.551 | 0.018 | 0.12 | 0.73 | 0.115 | 0.733 | 1.399 | 0.000 |
|  | *MEIS1* | -0.861 | 0.025 | -0.603 | 0.151 | 1.643 | 0 | 0.945 | 0.005 | 2.024 | 0.000 | 1.602 | 0.000 |
|  | *TCF7L2* | -0.673 | 0.465 | -0.5 | NA | -0.104 | 0.918 | 1.004 | 0.203 | 0.582 | 0.339 | 0.948 | 0.167 |
|  | *HES1* | 0.736 | 0.26 | 0.42 | NA | 1.132 | 0.009 | 0.17 | 0.889 | 0.227 | 0.632 | -0.101 | 0.910 |
|  | *NR4A1* | 1.166 | 0.012 | 0.301 | NA | 0.441 | 0.512 | 0.536 | 0.574 | 0.101 | 0.890 | 0.949 | 0.105 |
|  | *MECOM* | 0.173 | 0.956 | 0.129 | NA | -0.431 | 0.369 | 0.154 | 0.87 | -0.294 | 0.499 | -0.530 | 0.394 |
| RIBOSOME SIGNATURE | *ASCC3* | 0.328 | 0.494 | -0.018 | 0.958 | -0.379 | 0.149 | -0.729 | 0.000 | -0.476 | 0.041 | -0.348 | 0.013 |
|  | *HSPA14* | 0.035 | 0.991 | 0.233 | 0.398 | -0.369 | 0.009 | -0.569 | 0.015 | -0.393 | 0.003 | -0.508 | 0.007 |
|  | *MRPL15* | 0.172 | 0.871 | 0.334 | 0.019 | -0.380 | 0.079 | -0.594 | 0.000 | -0.544 | 0.004 | -0.877 | 0.000 |
|  | *MRPL27* | 0.111 | 0.930 | 0.224 | 0.117 | -0.270 | 0.138 | -0.368 | 0.003 | -0.406 | 0.009 | -0.599 | 0.000 |
|  | *MRPL3* | 0.080 | 0.968 | 0.190 | 0.133 | -0.127 | 0.596 | -0.504 | 0.000 | -0.240 | 0.172 | -0.472 | 0.000 |
|  | *MRPL4* | 0.234 | 0.641 | 0.367 | 0.001 | -0.271 | 0.246 | -0.329 | 0.009 | -0.580 | 0.001 | -0.856 | 0.000 |
|  | *MRPL58* | 0.176 | 0.762 | 0.192 | 0.322 | -0.244 | 0.225 | -0.425 | 0.011 | -0.416 | 0.010 | -0.555 | 0.000 |
|  | *MRPS17* | 0.047 | 0.990 | 0.120 | 0.423 | -0.243 | 0.213 | -0.502 | 0.000 | -0.360 | 0.028 | -0.602 | 0.000 |
|  | *MTG2* | 0.123 | 0.810 | 0.328 | 0.157 | -0.261 | 0.047 | 0.013 | 0.974 | -0.381 | 0.001 | -0.599 | 0.001 |
|  | *NSUN4* | 0.140 | 0.827 | 0.124 | 0.769 | -0.645 | 0.000 | -0.475 | 0.126 | -0.684 | 0.000 | -0.624 | 0.008 |
|  | *RPL22L1* | 0.175 | 0.797 | 0.144 | 0.283 | -0.586 | 0.000 | -1.028 | 0.000 | -0.865 | 0.000 | -1.019 | 0.000 |
|  | *RPL7L1* | 0.310 | 0.053 | 0.278 | 0.019 | -0.230 | 0.117 | -0.438 | 0.000 | -0.544 | 0.000 | -0.519 | 0.000 |

The DEGs between control vs sh*BCL11A* Erys without GC included higher expression of *CDKN2D* (both at day 3 and 5), a cell cycle inhibitor [28], and downregulation of *FOS* (at day 3) and upregulation of *MEIS1* (both day 5 and 3), and *HES1* (at day 3) [29] (**Table 1**). *CDKN2D* and *MEIS2* were altered also in sh*BCL11A* Erys with GC with respect to control. Furthermore, DEGs between control vs sh*BCL11A* Erys without and with GC included a strong downregulation of genes encoding ribosomal proteins (six genes without GC, *RPL22L1*, *MRPL3, MRPL15*, *MRPS17*, *HSPA14* and *SC3*; 10 genes with GC, *RPL7L1*, *RPL22L1*, *MRPL4*, *MRPL15*, *MRPL27*, *MRPL58*, *MRPS17*, *MTG2*, *NSUN4* and *HSP14*). The KEGG analyses identified that by day 5, sh*BCL11A* cells without GC expressed down-regulation of pathways involved in initiation of mRNA translation, ribosome biosynthesis, and cell cycle, while pathways linked to hematopoietic cell lineage, cytokine receptor interactions and JAK/STAT were upregulated (**Figure 6C**). Many of these pathways were also detected using the Reactome and Hallmark gene sets (**Figures 7,S6**). Of interest, however, the hallmark gene set analyses also identified that by day 5, sh*BCL11A* cells exhibited downregulation of genes involved in heme metabolism and upregulation of those involved in apoptosis (**Figure S7**).

**Figure 7.**
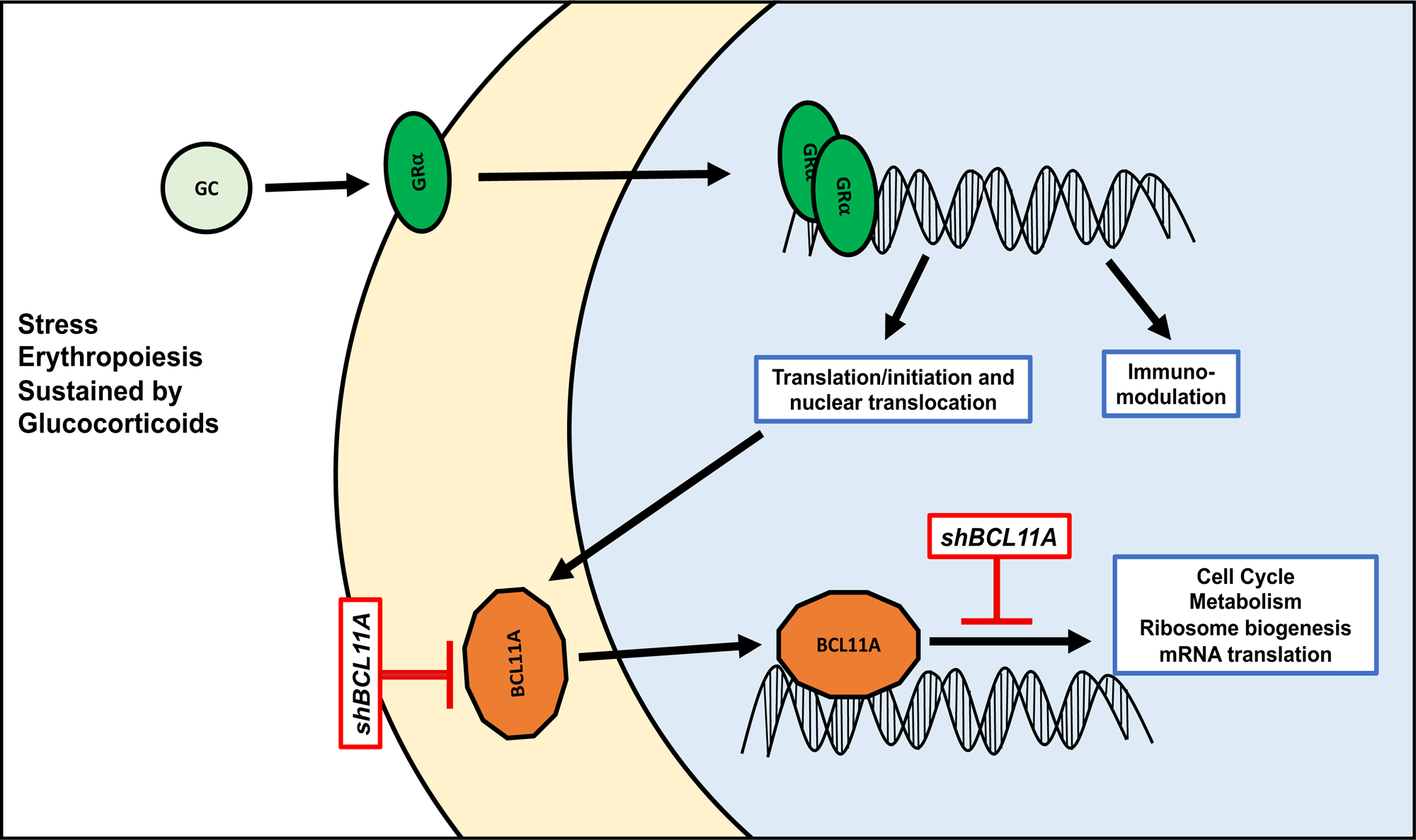
Summary of the proposed mechanism for the relationship between the glucocorticoid receptor and BCL11A in eliciting the erythroid response to glucocorticoids. We hypothesis that in erythroid cells, BCL11A, in addition to the globin genes, regulates the expression of genes involved in cell cycle, mRNA translation (ribosome biosynthesis) and metabolism (mitochondrial activity and lipid metabolism). Activation of the GC receptor induces early and late responses. The early responses include activation of translation initiation and nuclear translocation and is BCL11A independent. This early activity increases the protein content in the nucleus of BCL11A and leads to the late cellular response which highjacks and potentiate the activity of BCL11A in erythroid cells. This hypothesis provides a molecular mechanism for the reduced amplification in vitro in response to GC of cells with low BCL11A activity, either because they carry microdeletions of the genes or because transduced with shRNA and may have implications for the reduced response to GC of some DBA patients.

KEGG analyses also identified down-regulation of pathways involved in ribosome biosynthesis (both at day 3 and 5), cell cycle (day 3) in sh*BCL11A* cells with GC with respect to control while pathways linked to hematopoietic cell lineage (day 5) were upregulated (**Figure 6D**). Numerous additional pathways (such as P53 signaling, mismatch repair and DNA replication) were downregulated in sh*BCL11A* cells with GC at day 5. Many of these pathways were also detected using the Reactome and Hallmark gene sets (**Figures S6,7**). Of interest, the hallmark gene set analyses also identified that by day 5, sh*BCL11A* cells without GC, but not those with GC, exhibited downregulation of genes involved in heme metabolism and upregulation of those involved in apoptosis (**Figure S7**). Overall, this expression signature is consistent with the interpretation that the reduced expansion of sh*BCL11A* cells in the presence of GC is due to reduced proliferation and accelerated differentiation rather than apoptosis.

All together the above data indicate that *BCL11A* in addition to its silencing role of *HBG* regulates cell cycle related and other metabolic activities of Erys and these are compromised when its levels are reduced.

## DISCUSSION

*BCL11A* encodes a zinc finger protein and has been identified by genome-wide association studies (GWAS) [30,31] as the strongest quantitative trait locus yet discovered to explain variation in fetal globin (*HbF)* expression in patients with sickle cell anemia and β-thalassemia [31,32]. Established as a key repressor of *HbF* by multiple studies, its expression is inversely correlated with *HBG* levels in Erys generated both *in vitro* [22] and *in vivo* [27]. Most recent studies documented that BCL11A, along with the other developmental regulator ZBTB7A[33], directly represses the fetal *HBG* promoters by binding to well defined elements (*BCL11A* to - 115bp and *ZBTB7A* to -200bp) and recruitment of the suppressor NuRD complex [34].

The *BCL11A* gene itself is expressed as four major isoforms (XL, L, S and XS) [35]. The XL and L forms are reported to predominate in primary adult human Erys [27], while in embryonic and fetal cells (fetal liver and primitive erythroblasts or K562 cells that express mainly *HBG*), total *BCL11A* expression is lower and the shorter isoforms predominate [27,35].

In addition to the important role on the silencing of *HBG*, BCL11A has additional prominent effects. Alteration of the expression of *BCL11A* is associated with dysregulation of the hematopoietic system and progression to B cell malignancies[36]. Knockdown of *BCL11A* leads to quiescence, decreasing the proliferation of hematopoietic stem cells [29,37] while its overexpression is associated with the growth of some solid tumors, such as laryngeal squamous cell carcinoma and neuroblastoma [38,39]. In this study, we explored the cellular and molecular consequences of BCL11A on adult Erys in response to corticosteroids.

### GC effects on erythroid cells

We demonstrated that CD34+ cells from healthy donors cultured with GC contain greater levels of BCL11A, especially in the nucleus, suggesting that BCL11A is a novel factor in the GR signaling cascade.

To further investigate whether increases in BCL11A expression had functional significance, we cultured CD34+ cells from patients with *BCL11A*-microdeletions with or without GC. Patients’ Erys had decreased proliferative expansion after GC addition, but they had no difference from controls when GC was not added. Similarly, HUDEP-2 and normal CD34+ cells with enforced BCL11A reduction did not respond to GC-induced proliferation compared to control cells. These findings pinpoint a novel association between BCL11A and the proliferative response to GC of Erys.

### Molecular consequences of GC addition

The molecular details of the effects of GC on *BCL11A* expression is a work in progress. Previous data indicated that KLF1 is an upstream regulator of BCL11A activity during globin switching [40]. Later studies indicated that translation impairment is the major mechanism reducing BCL11A activity in fetal Erys [41]. We documented that Erys generated in the presence of Dex contain higher levels of *KLF1* (**Figure S8**). Furthermore, CD34+ cells from patients with congenital dyserythropoietic anemia due to the *KLF1*-E325K mutation express low levels of *BCL11A* (**Table S3**) and expand poorly *in vitro* in response to GC [42]. While expecting that the effects of GC on *BCL11A* expression may be at the transcriptional level and mediated through KLF1, we found that the level of mRNA for both *KLF1* and *BCL11A* in control cells plus and minus GC were similar (**Table 1**), displaying instead a strongly activated translation initiation and nuclear translocation signatures. These results suggest that GC may increase the activity of BCL11A (and of KLF1) indirectly by favoring the expression of genes which promote the translation of its mRNA and the nuclear translocation of its protein (**Figure 7**). Support of this hypothesis is provided by the fact that translation initiation and nuclear translocation pathways are also observed in control cells exposed to GC.

The reduced DEGs observed in shBCL11A cells treated with or without GC support the thesis that *BCL11A* is required to elicit a molecular response to GC in Erys. This is more striking in comparison with the robust changes in gene expression induced by GC in the control cells. The genes affected by GC in the control cells identify significant alterations in numerous pathways. Most notably, both at day 3 and day 5, GC activated robust mRNA translation and mitochondrial signatures, two pathways which impact erythroid proliferation and differentiation [43]. By contrast, the very few pathways enriched in sh*BCL11A* cells with and without GC at day 5, confirm that the response of BCL11A-depleted cells to GC is impaired.

Recently it has been demonstrated that Erys are important components of the innate immune response [44]. Single cell RNAseq has identified that fetal liver and cord blood contain a subset of Erys that express, like adult Erys generated in vitro with GC, high levels of CD36 which has an immune-modulatory signature and activates the immune functions of neutrophils and macrophages [45]. Given the well-known immune-modulatory role of GC, it is not surprising that a great number of the pathways regulated by GC in control cells belong to the immune lineage. Of interest, some of these immune pathways are also downregulated by GC in the sh*BCL11A* cells by day 5. These results suggest that the immune-modulatory functions of GC in Erys are mostly *BCL11A*-independent (**Figure 7**).

The greatest variability in gene expression was observed between control and sh*BCL11A* cells cultured with and without GC. In particular, the control and sh*BCL11A* treated cells cultured without GC expressed, in addition to *HBG*,changes in 1719 other genes (1090 upregulated, 629 downregulated) compared to those cultured with GC. This observation is in line with emerging data obtained from the Orkin laboratory which indicated that, in addition to regulating *HBG* expression [27], BCL11A is an important regulator of other functions of Erys [46]. Among the genes differentially expressed by control vs sh*BCL11A* treated cells we have identified significant downregulation of *CDKN2D,* a cell cycle inhibitor [28], and downregulation of *FOS* and upregulation of *MEIS1*, and *HES1*[29]. This expression profiling suggests that the trend toward reduced colony growth of CD34+ cells from the microdeletion patients with respect to their parents and the reduced baseline expansion of sh*BCL11A* cells with respect to control may highlight a role of *BCL11A* as cell cycle regulator of adult Erys independently from GC. Additional studies, outside the purpose of the current paper, coupled with loss of function of the putative *BCL11A*-target genes, are necessary to elucidate the full spectrum of *BCL11A* functions in erythropoiesis.

### GC and fetal HbF expression in vivo and in vitro

Stress erythropoiesis *in vitro* is characterized by an increase of HbF [5,6], but this has not been linked to changes in known globin regulators (BCL11A, KFL1, ZBTB7A) [47]. The *HBG*/*HBB* ratio is higher in immature than in mature cells because HbF production is first initiated in immature cells and then diluted out with maturation [5].

*In vivo* upregulation of HbF, following bleeding or post bone marrow transplantation, is also due to stress and is transient[48,49]. Furthermore, GC treatment did not change HbF levels *in vivo* in DBA patients with presumed stress erythropoiesis [50,51] and in patients with constitutive GR activation (Cushing’s) [16]. Thus, it appears that incremental changes in BCL11A above saturating levels do not further change HbF levels but do influence other important Ery functions. Most recently [52], it was shown that longest cell cycles sustained by *p27^Kip1^* (*CDKN1B*) raise the level of HbF (higher in slow cycling cells) by, at least in part, reducing *BCL11A*.This result is reminiscent of a previously formulated hypothesis that the in vitro increase of HbF is likely brought about by compression of differentiation/maturation kinetics[53]. Further studies are necessary to elucidate if indeed cell cycle acceleration is induced by BCL11A and whether *BCL11A*, in addition to its direct effects on *HbF* (by promoter binding), may induce *HbF* silencing also through changes in cell cycle.

In conclusion, our data indicate that GC exert both immune-modulatory and expansion activities in Erys and that its expansion activity is highly dependent on *BCL11A*.

## Acknowledgments

The authors wish to gratefully thank Drs. Paolo Prontera and Maria Piccione for providing blood from the microdeletion patients and their parents and Dr James Bieker for discussion.

## Authors Contribution

MM, JC, TN, LV, FM and AF performed experiments and analyzed the data. MM and JC performed the biostatistical analyses. MM, JC, TN, LV, FM and AF revised the manuscript. TP and ARM designed the study, interpreted the data and wrote the manuscript. All the authors read the manuscript and concur with its content.

