## Supplementary for "The glucocorticoid receptor elicited proliferative response in human erythropoiesis is BCL11A-dependent"

### SUPPLEMENTARY FILES

### MATERIALS AND METHODS

**Human subjects.** Blood from 25 normal donors (healthy donors, HD) was provided as de-identified material by the Italian Red Cross Blood Bank (Rome, Italy) according to guidelines established by local ethical human subject committees. CD34+ cells from de-identified healthy donors were purchased from the Fred Hutchison Cancer Center (Seattle, WA, USA). Blood from three heterozygous *BCL11A* microdeletion patients and from their parents was provided by the University of Perugia and Palermo (Italy) [1]. Patient 1 harbors a 3.5 Mb deletion downstream of the *BCL11A* gene, and while the *BCL11A* gene itself appears to be intact, its expression is considerably downregulated, potentially due to the absence of downstream regulatory elements required for normal expression[1]. Patients 2 and 3 carry 642 kb and 2.5 Mb deletions, respectively, that encompass the entire *BCL11A* open reading frame. Clinical details for these patients are previously provided[1].

**Proliferation (Prol) and differentiation (Diff) erythroid cultures.** Blood mononuclear blood cells were separated by centrifugation at 400gx30min over Ficoll-Hypaque. Human erythroblasts were obtained by culturing blood mononuclear cells ($10^6$ cells/mL) for 10-15 days in Iscove's modified Dulbecco's medium (IMDM) containing fetal bovine serum (FBS, 20%), SCF (10 ng/mL), EPO (1U/mL), interleukin-3 (IL-3, 1ng/mL), dexamethasone (Dex, $10^{-6}$ M) and estradiol ($10^{-6}$ M), as described[2]. Selected experiments were conducted in parallel cultures supplemented with growth factors plus and minus Dex. Day 8-12 proliferating erythroblasts (Prol) were collected and induced to differentiate (Diff) by culture for 4 additional days with EPO (3U/mL) and human recombinant insulin (10ng/mL).

**Colony forming cell (CFC) determinations.** Blood mononuclear cells ($3x10^5$ cells/mL) were cultured in MethoCult cultures stimulated with SCF (10ng/mL), IL-3 (10ng/mL), G-CSF (100ng/mL) and EPO (5U/ml), as described[3]. The cloning potential of adult CD34+ cells ($6x10^3$ cells/mL) and of their progeny expanded in culture was determined by culturing the cells in MethoCult™ H4434 Classic. The dishes were incubated at 37°C and 5% $pCO_2$ in a fully humidified incubator and colony growth scored at day 14. Colonies derived from BFU-E, CFU-GM, and CFU-GEMM were recognized according to standard morphological criteria[3].

**Morphological analyses.** Morphological observations were performed on cytocentrifuged smears (Shandon, Astmoor, UK) stained with May-Grünwald-Giemsa with the Axioscope light microscope equipped with a Coolsnap video camera (Zeiss, Oberkochen, Germany).

**RNA isolation and quantification of gene expression by real-time PCR.** Total RNA was isolated from $10^6$ PROL and DIFF cells using Trizol and reverse transcribed with the RNase OUT kit, as described by the manufacturers. Quantitative real-time PCR was carried out in a 7700 Sequence 14 Detection System (Applied Biosystems, Foster City, CA, USA), employing the TaqMan Master Mix containing AmpliTaq Gold DNA polymerase using amplification primers and probes already described[4]. *BCL11A* expression was measured according to the manufacturer's instructions with a kit which amplifies all three mRNA gene isoforms, as well as total mRNA (**Figure S1**). Each sample was analyzed in triplicate and the variability between these independent determinations was always within 5%. The mean of these determinations was used to calculate the mRNA level and expressed in arbitrary units, using the mean of the *hGAPDH* determinations as a calibrator, according to manufacturer's instructions. Expression levels were calculated with the algorithm ΔCt=CtX–Ct*GAPDH*, where Ct is the average threshold cycle and X is the gene being analyzed, and presented as $2^{-\Delta Ct}$. The expression ratio *HBG*/(*HBG*+*HBB*) was calculated as $2^{-\Delta Ct}$*HBG*/($2^{-\Delta Ct}$*HBG*+$2^{-\Delta Ct}$*HBB*). In selected experiments, PCR amplified fragments were separated by gel electrophoresis, eluted from the gel, purified with the QIAquick PCR purification kit and sequenced by the core facility of MSSM to verify that the amplification of each *BCL11A* isoform was correct. To asset the knock down of *BCL11A* in HUDEP-2 and CD34+ cells, RNA was extracted from $5x10^4$-$3x10^5$ cells using the Qiagen RNeasy Micro Kit. For *BCL11A* and globin determinations, RNA was reverse transcribed with the iScript Reverse Transcription Supermix and expression levels were calculated as relative expression with respect to levels of the housekeeping gene *B2M*, as described[5].

**Western Blot analysis.** Whole cell extracts (30μg protein/lane) were separated on SDS-PAGE and transferred to nitrocellulose membranes which were then probed with primary antibodies against BCL11A, GATA1, HBB, HBG, GRα, GAPDH, Lamin B1 and appropriate horseradish peroxidase-coupled secondary antibodies. For HUDEP-2 and CD34+ western blots, nuclear protein fractions (15μg protein/lane) were probed with antibodies against BCL11A and Histone 3 and with goat anti-mouse horseradish peroxidase-coupled secondary antibodies. Cells were subjected to sub-cellular fractionation with the NE-PER nuclear and cytosolic extraction kit, as described[6].

**Flow cytometry.** The erythroid differentiation of the progeny of CD34+ cells was assessed by incubating the cells for 30min at 4°C in the dark with Phycoerythrin (PE)-CD34, Pacific Blue 450 (PB450)-CD36 and fluorescein isothiocyanate (FITC)-CD235a. The cells were washed twice with PBS and antibody binding determined with the Beckman Coulter FACS. Data were analyzed with FlowJo.

**RNA-Seq for *BCL11A* microdeletion patients.** RNA was extracted from $1x10^6$ cells using TRIzol, cleaned with miRNeasy Mini kit and used to prepare RNA libraries. Sequencing, data processing and alignment were performed as described[1] .

**Culture of HUDEP-1 and HUDEP-2 cells and knock down of *BCL11A*.** Cell lines were maintained in StemSpan SFEM, containing penicillin/streptomycin (2%), L-Glutamine (1%), SCF (100µg/mL), EPO (100U/mL), and doxycycline (100µg/mL) with or without Dex ($10^{-6}$mg/mL), as described[7]. Cultures were incubated at 37°C and 5% $pCO_2$ in a fully humidified incubator. HUDEP-2 cells were transduced ($1x10^5$ cells/0.4µL of virus) either with control shRNA (shRNA against Luciferase, with U6 promoter driving the shRNA and the *hPGK* promoter driving the puromycin resistance gene, *shLuc*) or the *BCL11A* shRNA (*shBCL11A*) in the presence of synperonic (0.25mg/mL).

**Knock down of *BCL11A* in CD34+ cells.** CD34+ cells were transduced with either *shLuc* or sh*BCL11A*, as described[8] (**FigureS2)**. Briefly, CD34+ cells were thawed, cultured for 2 days in expansion media (StemSpan H3000 media with CC110 supplements, penicillin/streptomycin, small molecules (SR1, Ly, UM171, 1000x) and protamine sulfate (8ug/mL)). On Day -5, CD34+ cells were transduced ($1x10^5$ cells/3.2µL of virus) with the same viruses used for HUDEP-2 cells. Before transduction, the 24 wells plates had been coated with 50 ug/mL retronectin. The transduced CD34+ cells were cultured for 3 additional days in expansion media plus puromycin (0.5µg/mL). At Day 0, the cells ($2x10^5$ cells/mL) were transferred in erythroid expansion media composed by IMDM, human adult plasma (5%), heparin (2IU/mL), insulin (10µg/mL), transferrin (330 µg/mL), SCF (100ng/mL), EPO (3U/mL), IL-3 (5ng/mL), plus and minus hydrocortisone (1µM). These knock-down experiments were performed with hydrocortisone because preliminary experiments had determined that this GR agonist is equally potent to Dex in expanding human CD34+ cells (**Figure S3**) and for consistency with transduction protocols published in earlier studies[8]. Since GC exerts their effects mostly until day 7[9,10], cells were cultured without hydrocortisone from day 7 on, as reported[8,10]. Day 0

CD34+ cells were characterized by determining CFC content and *BCL11A, HBG* and *HBB* mRNA levels (by quantitative RT-PCR) and BCL11A content by Western Blot.

**RNA-seq of the progeny of transduced CD34+ cells**. The progeny of the shRNA control and shRNA *BCL11A* CD34+ cells ($5x10^4$-$3x10^5$) obtained with or without GC were harvested at day 3 and day 5 of culture for RNA-seq analysis. Illumina libraries were constructed using Alt-Seq V3 kit derived from prime-seq[11] and sequenced at read lengths 28x8x94 on the Illumina Novaseq 6000 (Illumina). Mapping, demultiplexing and quantification of RNA-seq data were performed by STARsolo. GENCODE v25 was used for annotation. Gene counts were obtained using featureCounts and FPKM per gene were calculated using Cufflinks. Differentially expressed genes (DEGs) between the different groups were analyzed with R package DESeq2.

**GeneSet Enrichment Analysis**. GSEA was performed using the GSEA software (GSEA 4.3.1 with default parameters). Pathway analyses were performed with the KEGG gene-set (c2.cp.kegg.v2023.1.Hs.symbols.gmt), REACTOME gene-set (c2.cp.reactome.v2023.1.Hs.symbols.gmt) and HALLMARK gene-set (h.all.v2023.1.Hs.symbols.gmt) downloaded from MSigDB (https://www.broadinstitute.org/gsea/msigdb/collections.jsp). Genes were pre-ranked in order of their differential expression between CD34+ treated or not treated with GC.

**Statistical Analyses.** Results are presented either as median, 25% to 75% interquartile range and maximum and minimum or as Mean (+/-SD), as most appropriate. Paired comparisons between *BCL11A* mRNA levels measured in Prol and Diff cells were obtained using the Wilcoxon signed-rank test. Values observed between cells treated with and without GC were compared by paired t-test. Statistical analysis of the experiments with HUDEP and CD34+ cells was performed with Anova multiple test. All statistical analyses were performed with the Graph Pad9 Prism software.

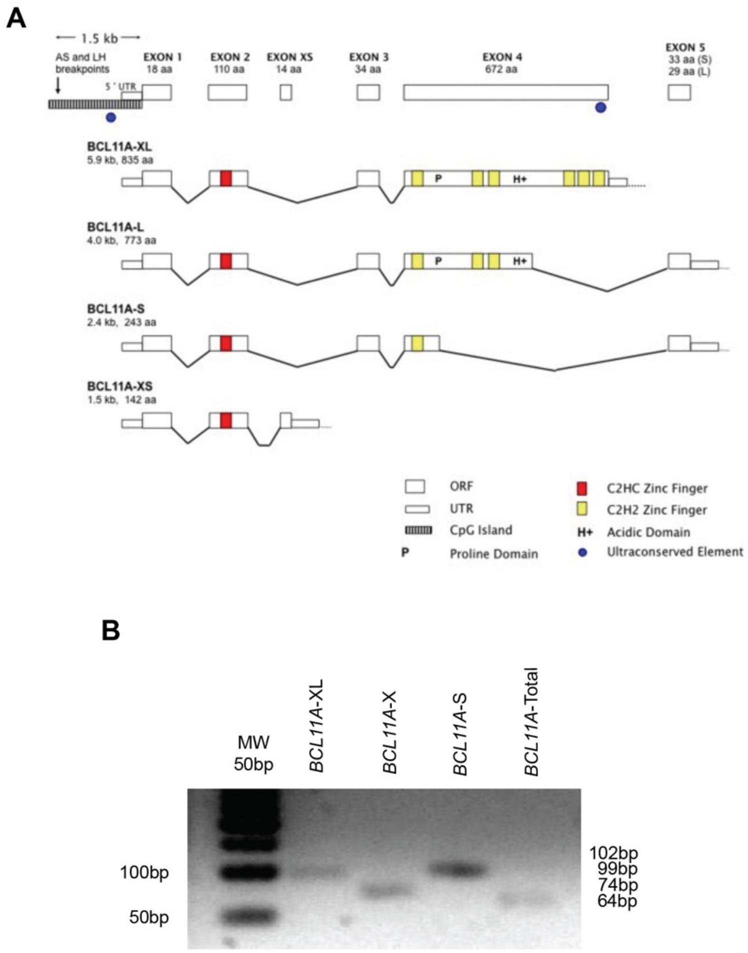


**Figure S1. Validation of the quantitative RT-PCR for BCL11A mRNA presented in Figure 1B.** A) The four major isoforms of BCL11A: extra-long (XL), long (L), short (S) and extra-short (XS)[12] and the position of the primers used to detect their expression by quantitative RT-PCR described in **Figure 1**B. The identity of the representative transcripts amplified by the various primers was confirmed by gel electrophoresis B) and sequencing (not shown).

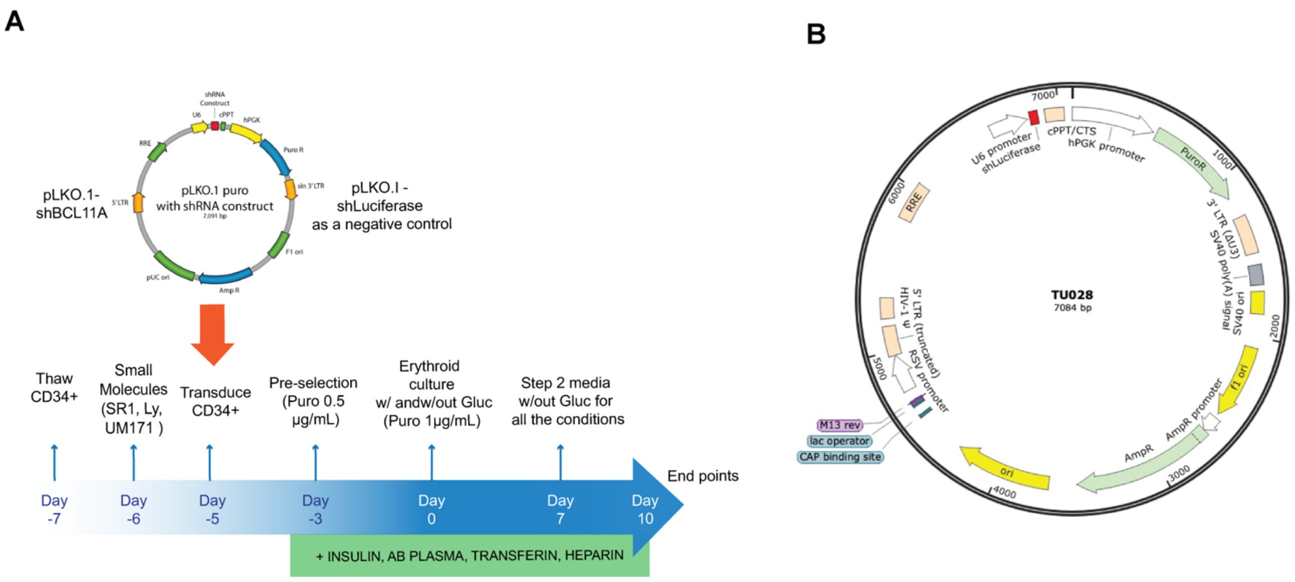


**Figure S2. Design of the experiments to test the effects of BCL11A loss of function of the response of adult CD34+ cells to glucocorticoids.** A) CD34+ cells were thawed and seeded at 2x10$^5$ cells/mL in StemSpan H3000 with CC110. After 24 hours, the smalls molecules (LY, UM171, SR1) were added to the culture media. The following day, CD34+ cells were transduced with either the *shBCL11A* (A) or the control *shLuciferase* vector (B) (3.2 µL/1x10$^5$ cells). The vectors contained the shRNA driven by the U6 promoter and the puromycin-resistance gene driven by the *hPGK* promoter. The transduced CD34+ were selected then with Puromycin (0.5 µg/mL) for three days and then cultured in erythroid expansion cultures for 7 days with and without GC, as described in[8,10]. On day 0, the CD34+ cells were characterized for their clonogenic potential and expression of *BCL11A* (both mRNA and protein). On day 3, 5, 7 and 10 of the erythroid expansion culture, the cells were counted and profiled for differentiation markers by FACS determinations with CD34, CD36 and GlyA(CD235a) and RNA-seq approach.

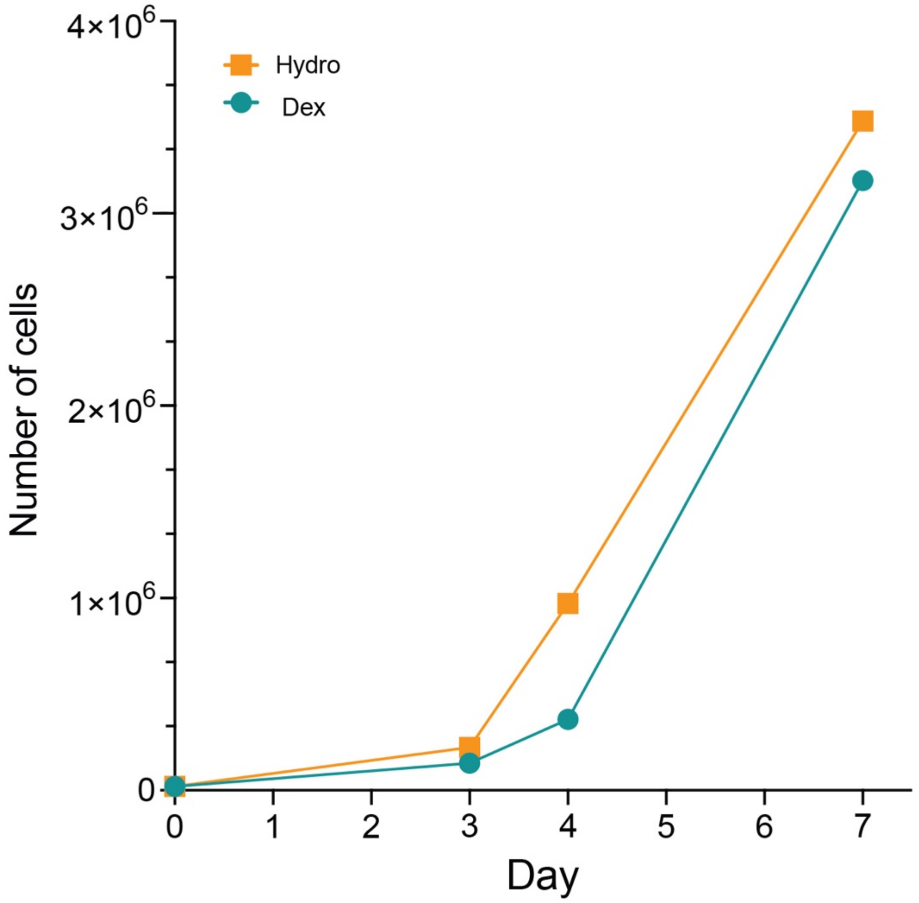


**Figure S3. Equivalent number of cells are generated over time by adult CD34+ cells when cultured in erythroid amplification media stimulated with Dexamethasone (Dex) or Hydrocortisone (Hydro).**

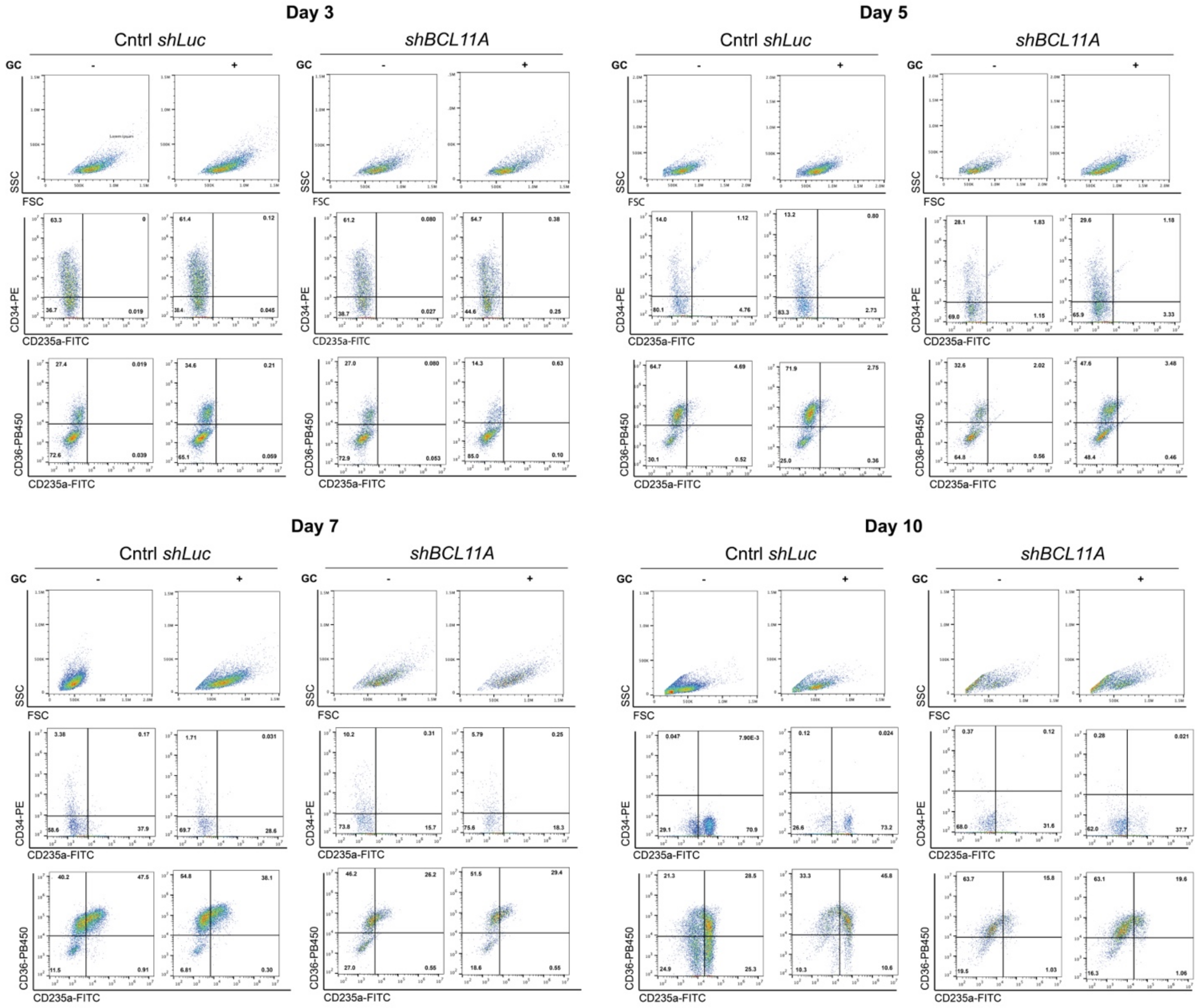


**Figure S4. Maturation profiling of the progeny of *shLuc* and *shBCL11A* CD34+ cells cultured in erythroid expansion media with or without GC for 3, 5, 7 and 10 days.** The presence of hematopoietic progenitors was monitored cells by profiling with CD34 and CD235a while the progression of stress-differentiation was monitored with CD36 and CD235a.

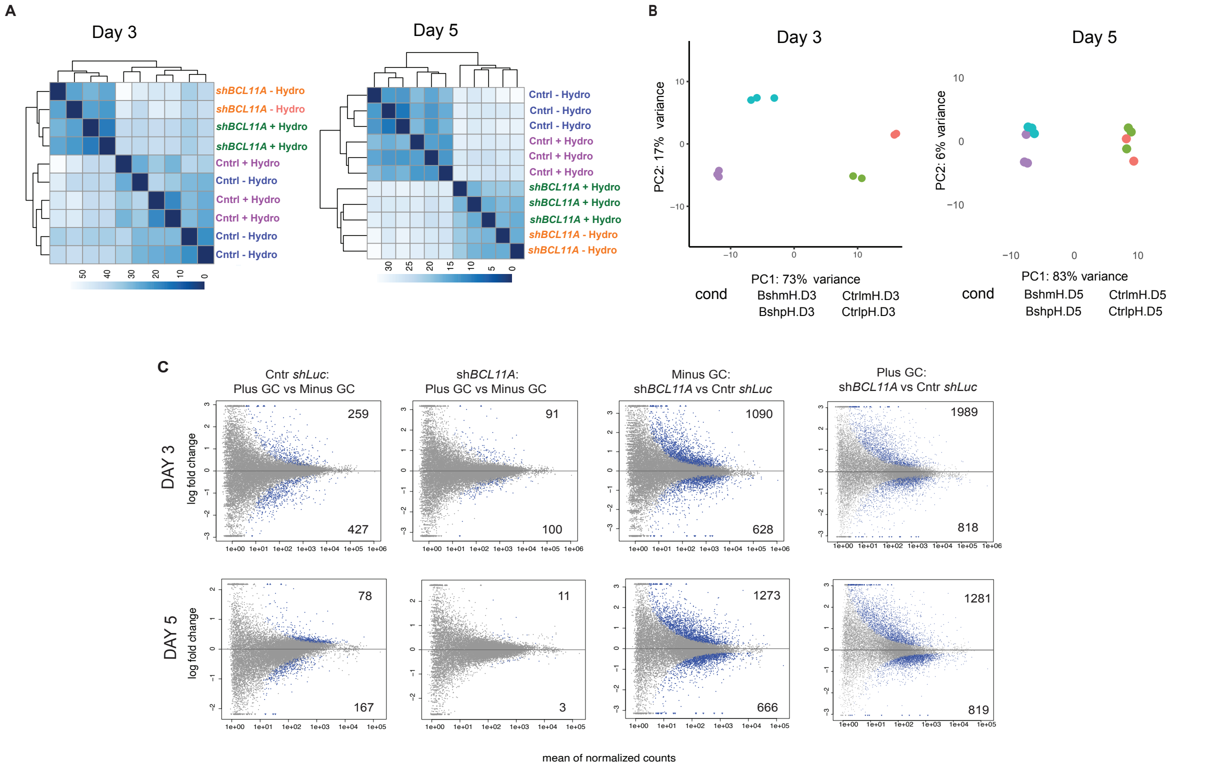


**Figure S5. Expression profiling of the progeny of control *shLuc* and sh*BCL11A* CD34+ cells cultured with or without GC for 3 and 5 days.** A) Heatmap showing the clusterization of the samples at day 3 and 5. **B)** Principal component analysis (PCA**)** of the RNAseq data obtained with the progeny of CD34+ cells transduced with control *shLuc* and sh*BCL11A* in presence or absence of GC at day 3 and day 5. **C**) MA plot indicating the number of genes expressed at significantly different levels between control *shLuc* cells cultured with and without GC, *shBCL11A* cells cultured plus and minus GC and between sh*BCL11A* and shLuc cells minus and plus GC at day 3 and 5 of culture.

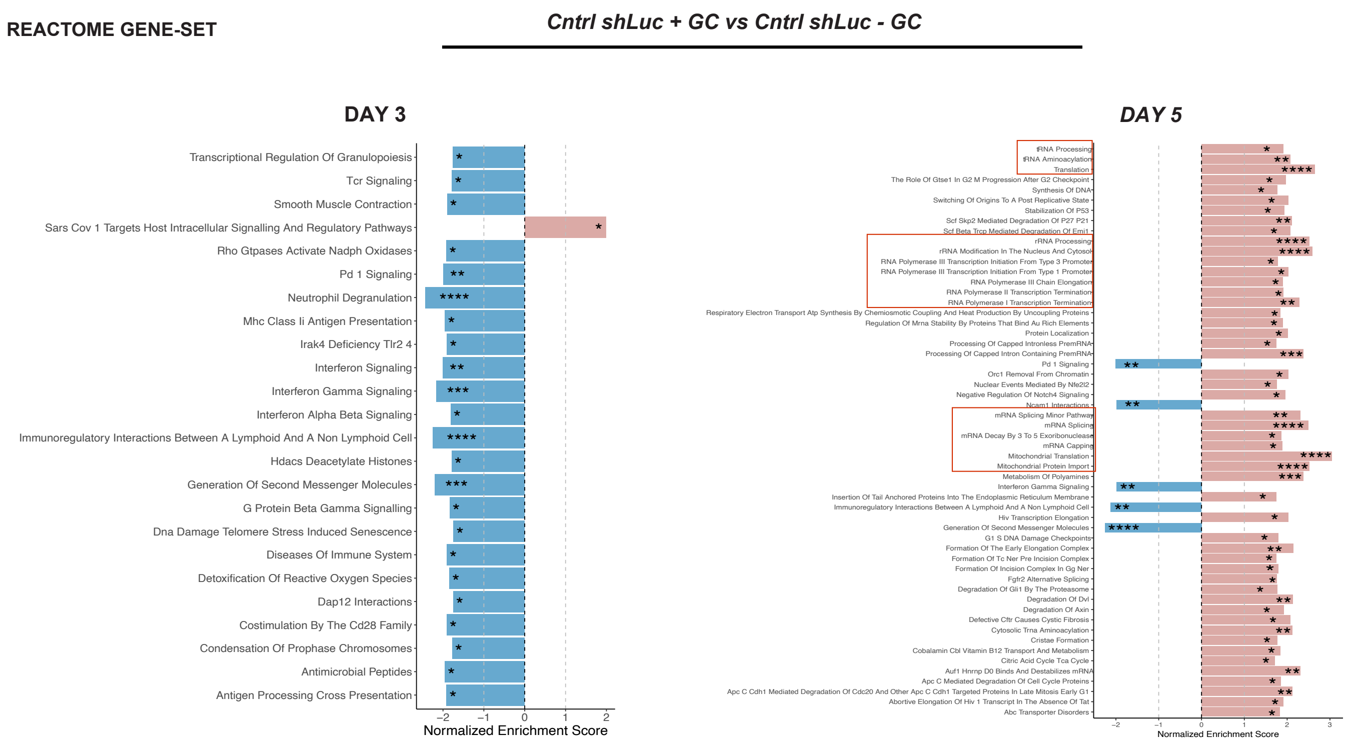
REACTOME GENE-SET
Cntrl shLuc + GC vs Cntrl shLuc - GC
DAY 3
Transcriptional Regulation Of Granulopoiesis
Tcr Signaling
Smooth Muscle Contraction
Sars Cov 1 Targets Host Intracellular Signalling And Regulatory Pathways
Rho Gtpases Activate Nadph Oxidases
Pd 1 Signaling
Neutrophil Degranulation
Mhc Class Ii Antigen Presentation
Irak4 Deficiency Tlr2 4
Interferon Signaling
Interferon Gamma Signaling
Interferon Alpha Beta Signaling
Immunoregulatory Interactions Between A Lymphoid And A Non Lymphoid Cell
Hdacs Deacetylate Histones
Generation Of Second Messenger Molecules
G Protein Beta Gamma Signalling
Dna Damage Telomere Stress Induced Senescence
Diseases Of Immune System
Detoxification Of Reactive Oxygen Species
Dap12 Interactions
Costimulation By The Cd28 Family
Condensation Of Prophase Chromosomes
Antimicrobial Peptides
Antigen Processing Cross Presentation
Normalized Enrichment Score
DAY 5
tRNA Processing
tRNA Aminoacylation
Translation
The Role Of Gtse1 In G2 M Progression After G2 Checkpoint
Synthesis Of DNA
Switching Of Origins To A Post Replicative State
Stabilization Of P53
Scf Skp2 Mediated Degradation Of P27 P21
Scf Beta Trcp Mediated Degradation Of Emi1
rRNA Processing
rRNA Modification In The Nucleus And Cytosol
RNA Polymerase III Transcription Initiation From Type 3 Promoter
RNA Polymerase III Transcription Initiation From Type 1 Promoter
RNA Polymerase III Chain Elongation
RNA Polymerase II Transcription Termination
RNA Polymerase I Transcription Termination
Respiratory Electron Transport Atp Synthesis By Chemiosmotic Coupling And Heat Production By Uncoupling Proteins
Regulation Of Mrna Stability By Proteins That Bind Au Rich Elements
Protein Localization
Processing Of Capped Intronless PremRNA
Processing Of Capped Intron Containing PremRNA
Pd 1 Signaling
Orc1 Removal From Chromatin
Nuclear Events Mediated By Nfe2l2
Negative Regulation Of Notch4 Signaling
Ncam1 Interactions
mRNA Splicing Minor Pathway
mRNA Splicing
mRNA Decay By 3 To 5 Exoribonuclease
mRNA Capping
Mitochondrial Translation
Mitochondrial Protein Import
Metabolism Of Polyamines
Interferon Gamma Signaling
Insertion Of Tail Anchored Proteins Into The Endoplasmic Reticulum Membrane
Immunoregulatory Interactions Between A Lymphoid And A Non Lymphoid Cell
Hiv Transcription Elongation
Generation Of Second Messenger Molecules
G1 S DNA Damage Checkpoints
Formation Of The Early Elongation Complex
Formation Of Tc Ner Pre Incision Complex
Formation Of Incision Complex In Gg Ner
Fgfr2 Alternative Splicing
Degradation Of Gli1 By The Proteasome
Degradation Of Dvl
Degradation Of Axin
Defective Cftr Causes Cystic Fibrosis
Cytosolic Trna Aminoacylation
Cristae Formation
Cobalamin Cbl Vitamin B12 Transport And Metabolism
Citric Acid Cycle Tca Cycle
Auf1 Hnrnp D0 Binds And Destabilizes mRNA
Apc C Mediated Degradation Of Cell Cycle Proteins
Apc C Cdh1 Mediated Degradation Of Cdc20 And Other Apc C Cdh1 Targeted Proteins In Late Mitosis Early G1
Abortive Elongation Of Hiv 1 Transcript In The Absence Of Tat
Abc Transporter Disorders
Normalized Enrichment Score

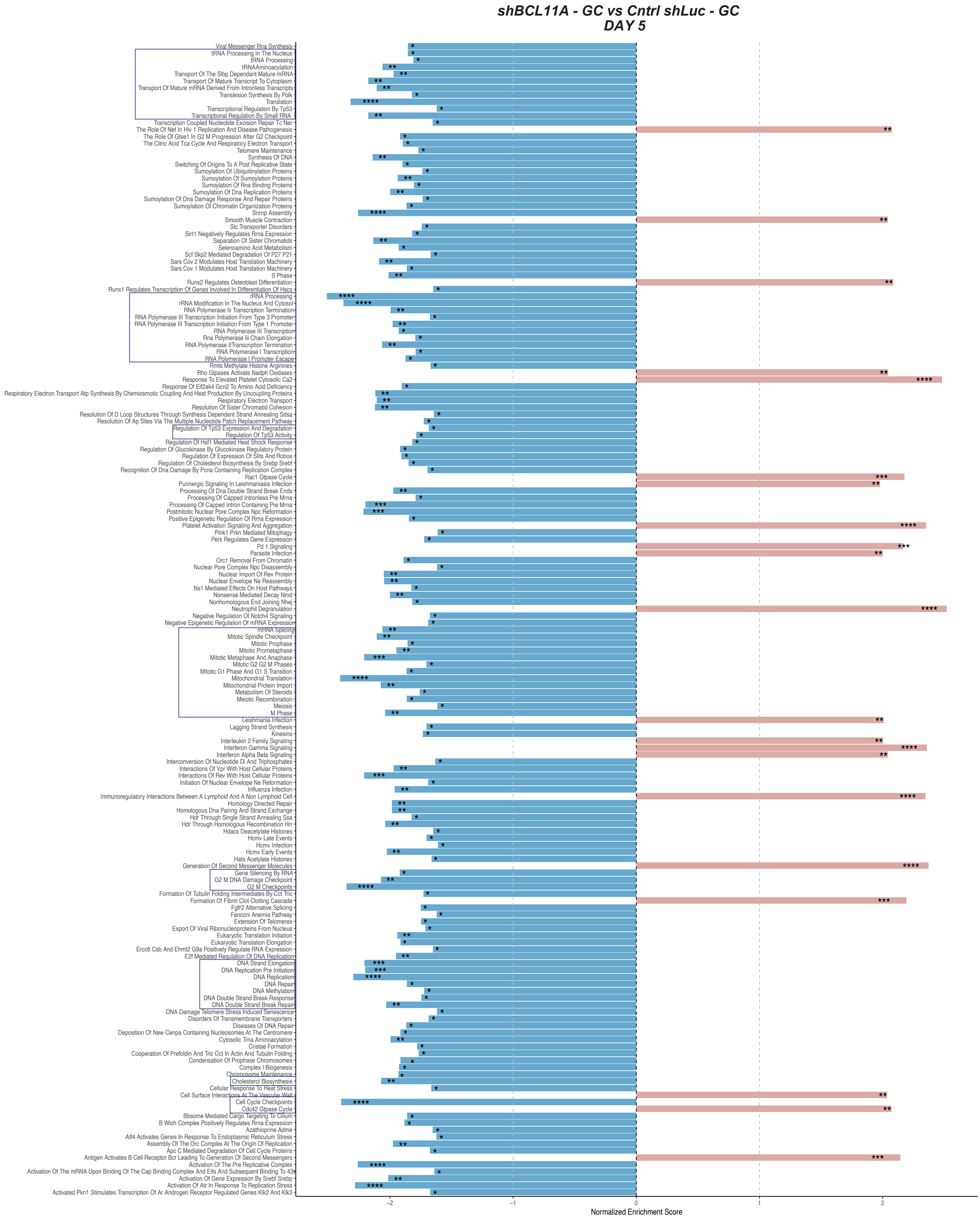
shBCL11A - GC vs Cntrl shLuc - GC
DAY 5
Viral Messenger Rna Synthesis
tRNA Processing In The Nucleus
tRNA Processing
tRNAAminoacylation
Transport Of The Slbp Dependant Mature mRNA
Transport Of Mature Transcript To Cytoplasm
Transport Of Mature mRNA Derived From Intronless Transcripts
Translesion Synthesis By Polk
Translation
Transcriptional Regulation By Tp53
Transcriptional Regulation By Small RNA
Transcription Coupled Nucleotide Excision Repair Tc Ner
The Role Of Nef In Hiv 1 Replication And Disease Pathogenesis
The Role Of Gtse1 In G2 M Progression After G2 Checkpoint
The Citric Acid Tca Cycle And Respiratory Electron Transport
Telomere Maintenance
Synthesis Of DNA
Switching Of Origins To A Post Replicative State
Sumoylation Of Ubiquitinylation Proteins
Sumoylation Of Sumoylation Proteins
Sumoylation Of Rna Binding Proteins
Sumoylation Of Dna Replication Proteins
Sumoylation Of Dna Damage Response And Repair Proteins
Sumoylation Of Chromatin Organization Proteins
Snrnp Assembly
Smooth Muscle Contraction
Slc Transporter Disorders
Sirt1 Negatively Regulates Rrna Expression
Separation Of Sister Chromatids
Selenoamino Acid Metabolism
Scf Skp2 Mediated Degradation Of P27 P21
Sars Cov 2 Modulates Host Translation Machinery
Sars Cov 1 Modulates Host Translation Machinery
S Phase
Runx2 Regulates Osteoblast Differentiation
Runx1 Regulates Transcription Of Genes Involved In Differentiation Of Hscs
rRNA Processing
rRNA Modification In The Nucleus And Cytosol
RNA Polymerase Iii Transcription Termination
RNA Polymerase III Transcription Initiation From Type 3 Promoter
RNA Polymerase III Transcription Initiation From Type 1 Promoter
RNA Polymerase III Transcription
Rna Polymerase Iii Chain Elongation
RNA Polymerase IITranscription Termination
RNA Polymerase I Transcription
RNA Polymerase I Promoter Escape
Rmts Methylate Histone Arginines
Rho Gtpases Activate Nadph Oxidases
Response To Elevated Platelet Cytosolic Ca2
Response Of Eif2ak4 Gcn2 To Amino Acid Deficiency
Respiratory Electron Transport Atp Synthesis By Chemiosmotic Coupling And Heat Production By Uncoupling Proteins
Respiratory Electron Transport
Resolution Of Sister Chromatid Cohesion
Resolution Of D Loop Structures Through Synthesis Dependent Strand Annealing Sdsa
Resolution Of Ap Sites Via The Multiple Nucleotide Patch Replacement Pathway
Regulation Of Tp53 Expression And Degradation
Regulation Of Tp53 Activity
Regulation Of Hsf1 Mediated Heat Shock Response
Regulation Of Glucokinase By Glucokinase Regulatory Protein
Regulation Of Expression Of Slits And Robos
Regulation Of Cholesterol Biosynthesis By Srebp Srebf
Recognition Of Dna Damage By Pcna Containing Replication Complex
Rac1 Gtpase Cycle
Purinergic Signaling In Leishmaniasis Infection
Processing Of Dna Double Strand Break Ends
Processing Of Capped Intronless Pre Mrna
Processing Of Capped Intron Containing Pre Mrna
Postmitotic Nuclear Pore Complex Npc Reformation
Positive Epigenetic Regulation Of Rrna Expression
Platelet Activation Signaling And Aggregation
Pink1 Prkn Mediated Mitophagy
Perk Regulates Gene Expression
Pd 1 Signaling
Parasite Infection
Orc1 Removal From Chromatin
Nuclear Pore Complex Npc Disassembly
Nuclear Import Of Rev Protein
Nuclear Envelope Ne Reassembly
Ns1 Mediated Effects On Host Pathways
Nonsense Mediated Decay Nmd
Nonhomologous End Joining Nhej
Neutrophil Degranulation
Negative Regulation Of Notch4 Signaling
Negative Epigenetic Regulation Of rRNA Expression
mRNA Splicing
Mitotic Spindle Checkpoint
Mitotic Prophase
Mitotic Prometaphase
Mitotic Metaphase And Anaphase
Mitotic G2 G2 M Phases
Mitotic G1 Phase And G1 S Transition
Mitochondrial Translation
Mitochondrial Protein Import
Metabolism Of Steroids
Meiotic Recombination
Meiosis
M Phase
Leishmania Infection
Lagging Strand Synthesis
Kinesins
Interleukin 2 Family Signaling
Interferon Gamma Signaling
Interferon Alpha Beta Signaling
Interconversion Of Nucleotide Di And Triphosphates
Interactions Of Vpr With Host Cellular Proteins
Interactions Of Rev With Host Cellular Proteins
Initiation Of Nuclear Envelope Ne Reformation
Influenza Infection
Immunoregulatory Interactions Between A Lymphoid And A Non Lymphoid Cell
Homology Directed Repair
Homologous Dna Pairing And Strand Exchange
Hdr Through Single Strand Annealing Ssa
Hdr Through Homologous Recombination Hrr
Hdacs Deacetylate Histones
Hcmv Late Events
Hcmv Infection
Hcmv Early Events
Hats Acetylate Histones
Generation Of Second Messenger Molecules
Gene Silencing By RNA
G2 M DNA Damage Checkpoint
G2 M Checkpoints
Formation Of Tubulin Folding Intermediates By Cct Tric
Formation Of Fibrin Clot Clotting Cascade
Fgfr2 Alternative Splicing
Fanconi Anemia Pathway
Extension Of Telomeres
Export Of Viral Ribonucleoproteins From Nucleus
Eukaryotic Translation Initiation
Eukaryotic Translation Elongation
Ercc6 Csb And Ehmt2 G9a Positively Regulate RNA Expression
E2f Mediated Regulation Of DNA Replication
DNA Strand Elongation
DNA Replication Pre Initiation
DNA Replication
DNA Repair
DNA Methylation
DNA Double Strand Break Response
DNA Double Strand Break Repair
DNA Damage Telomere Stress Induced Senescence
Disorders Of Transmembrane Transporters
Diseases Of DNA Repair
Deposition Of New Cenpa Containing Nucleosomes At The Centromere
Cytosolic Trna Aminoacylation
Cristae Formation
Cooperation Of Prefoldin And Tric Cct In Actin And Tubulin Folding
Condensation Of Prophase Chromosomes
Complex I Biogenesis
Chromosome Maintenance
Cholesterol Biosynthesis
Cellular Response To Heat Stress
Cell Surface Interactions At The Vascular Wall
Cell Cycle Checkpoints
Cdc42 Gtpase Cycle
Bbsome Mediated Cargo Targeting To Cilium
B Wich Complex Positively Regulates Rrna Expression
Azathioprine Adme
Atf4 Activates Genes In Response To Endoplasmic Reticulum Stress
Assembly Of The Orc Complex At The Origin Of Replication
Apc C Mediated Degradation Of Cell Cycle Proteins
Antigen Activates B Cell Receptor Bcr Leading To Generation Of Second Messengers
Activation Of The Pre Replicative Complex
Activation Of The mRNA Upon Binding Of The Cap Binding Complex And Eifs And Subsequent Binding To 43s
Activation Of Gene Expression By Srebf Srebp
Activation Of Atr In Response To Replication Stress
Activated Pkn1 Stimulates Transcription Of Ar Androgen Receptor Regulated Genes Klk2 And Klk3
−2
−1
0
1
2
Normalized Enrichment Score

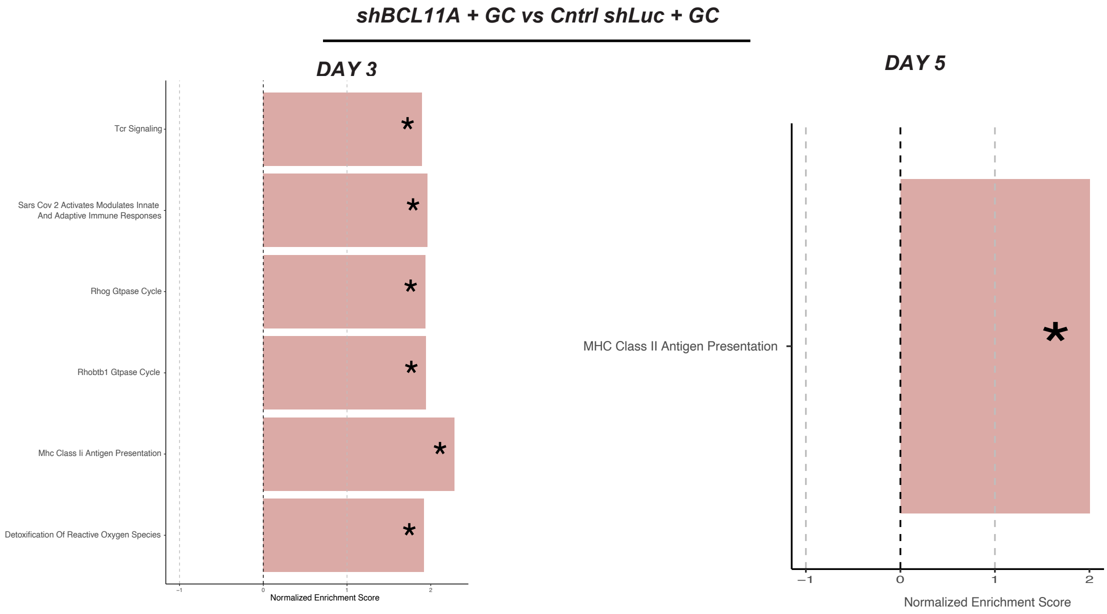


**Figure S6: GSEA enriched pathways analysis.** Comparison between control *shLuc* cells with or without GC at day 3 and day 5, between *shBCL11A* and control without GC at day 5 and between *shBCL11A* and control with GC at day 3 and 5. Red and blue columns indicate enriched and un-enriched pathway, respectively. All the data were compared using the REACTOME gene set (c2.cp.reactome.v2023.1.Hs.symbols.gmt). Red and blue boxes indicate respectively up- and down-enriched pathways. FDR q-values among groups are indicated by asterisks (*, p<0.05; **,p<0.001; ***; p<0.0001****; p<0.00001).

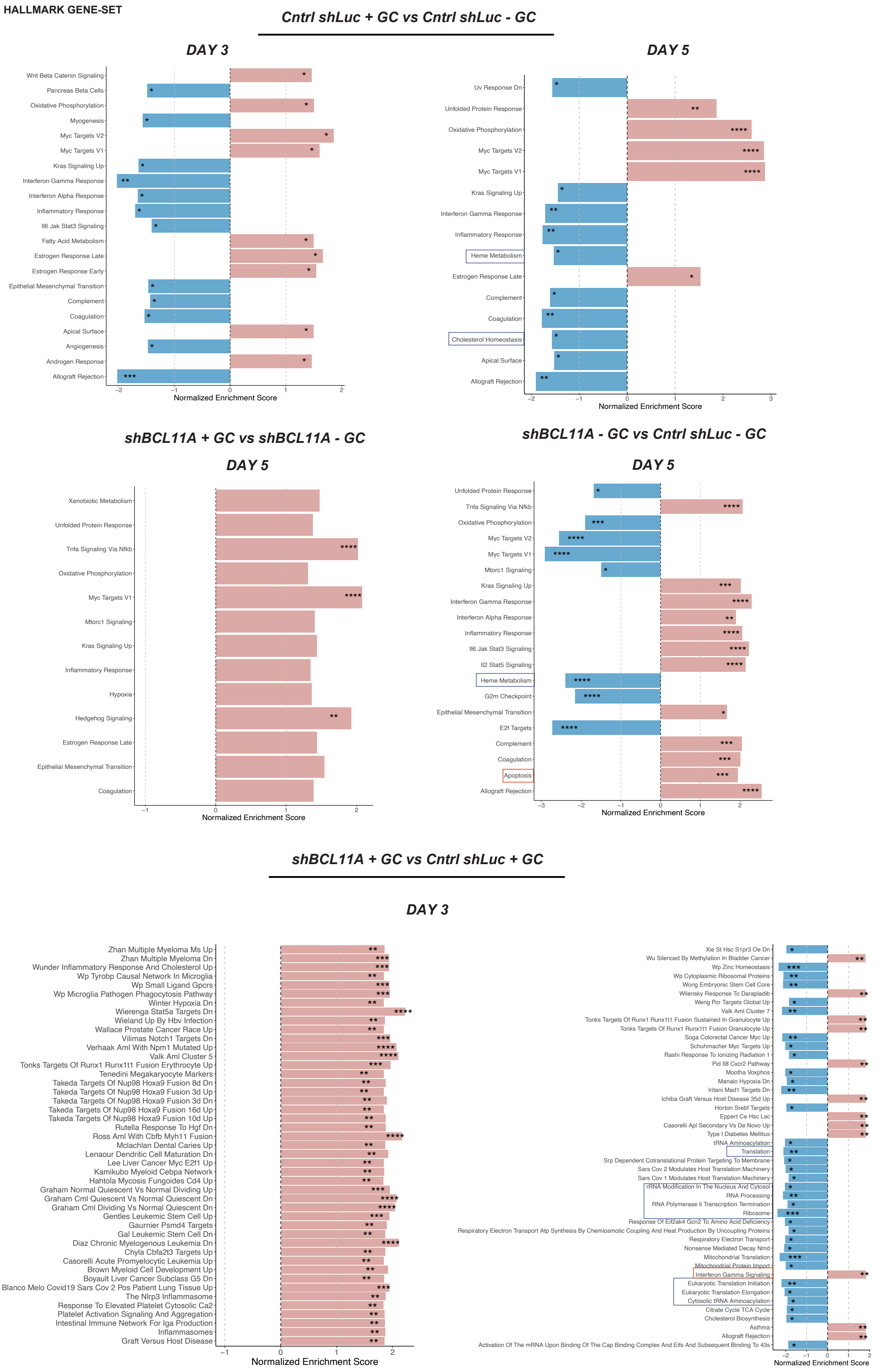

HALLMARK GENE-SET
Cntrl shLuc + GC vs Cntrl shLuc - GC
DAY 3
Wnt Beta Catenin Signaling
Pancreas Beta Cells
Oxidative Phosphorylation
Myogenesis
Myc Targets V2
Myc Targets V1
Kras Signaling Up
Interferon Gamma Response
Interferon Alpha Response
Inflammatory Response
Il6 Jak Stat3 Signaling
Fatty Acid Metabolism
Estrogen Response Late
Estrogen Response Early
Epithelial Mesenchymal Transition
Complement
Coagulation
Apical Surface
Angiogenesis
Androgen Response
Allograft Rejection
Normalized Enrichment Score
DAY 5
Uv Response Dn
Unfolded Protein Response
Oxidative Phosphorylation
Myc Targets V2
Myc Targets V1
Kras Signaling Up
Interferon Gamma Response
Inflammatory Response
Heme Metabolism
Estrogen Response Late
Complement
Coagulation
Cholesterol Homeostasis
Apical Surface
Allograft Rejection
Normalized Enrichment Score
shBCL11A + GC vs shBCL11A - GC
DAY 5
Xenobiotic Metabolism
Unfolded Protein Response
Tnfa Signaling Via Nfkb
Oxidative Phosphorylation
Myc Targets V1
Mtorc1 Signaling
Kras Signaling Up
Inflammatory Response
Hypoxia
Hedgehog Signaling
Estrogen Response Late
Epithelial Mesenchymal Transition
Coagulation
Normalized Enrichment Score
shBCL11A - GC vs Cntrl shLuc - GC
DAY 5
Unfolded Protein Response
Tnfa Signaling Via Nfkb
Oxidative Phosphorylation
Myc Targets V2
Myc Targets V1
Mtorc1 Signaling
Kras Signaling Up
Interferon Gamma Response
Interferon Alpha Response
Inflammatory Response
Il6 Jak Stat3 Signaling
Il2 Stat5 Signaling
Heme Metabolism
G2m Checkpoint
Epithelial Mesenchymal Transition
E2f Targets
Complement
Coagulation
Apoptosis
Allograft Rejection
Normalized Enrichment Score
shBCL11A + GC vs Cntrl shLuc + GC
DAY 3
Zhan Multiple Myeloma Ms Up
Zhan Multiple Myeloma Dn
Wunder Inflammatory Response And Cholesterol Up
Wp Tyrobp Causal Network In Microglia
Wp Small Ligand Gpcrs
Wp Microglia Pathogen Phagocytosis Pathway
Winter Hypoxia Dn
Wierenga Stat5a Targets Dn
Wieland Up By Hbv Infection
Wallace Prostate Cancer Race Up
Vilimas Notch1 Targets Dn
Verhaak Aml With Npm1 Mutated Up
Valk Aml Cluster 5
Tonks Targets Of Runx1 Runx1t1 Fusion Erythrocyte Up
Tenedini Megakaryocyte Markers
Takeda Targets Of Nup98 Hoxa9 Fusion 8d Dn
Takeda Targets Of Nup98 Hoxa9 Fusion 3d Up
Takeda Targets Of Nup98 Hoxa9 Fusion 3d Dn
Takeda Targets Of Nup98 Hoxa9 Fusion 16d Up
Takeda Targets Of Nup98 Hoxa9 Fusion 10d Up
Rutella Response To Hgf Dn
Ross Aml With Cbfb Myh11 Fusion
Mclachlan Dental Caries Up
Lenaour Dendritic Cell Maturation Dn
Lee Liver Cancer Myc E2f1 Up
Kamikubo Myeloid Cebpa Network
Hahtola Mycosis Fungoides Cd4 Up
Graham Normal Quiescent Vs Normal Dividing Up
Graham Cml Quiescent Vs Normal Quiescent Dn
Graham Cml Dividing Vs Normal Quiescent Dn
Gentles Leukemic Stem Cell Up
Gaurnier Psmd4 Targets
Gal Leukemic Stem Cell Dn
Diaz Chronic Myelogenous Leukemia Dn
Chyla Cbfa2t3 Targets Up
Casorelli Acute Promyelocytic Leukemia Up
Brown Myeloid Cell Development Up
Boyault Liver Cancer Subclass G5 Dn
Blanco Melo Covid19 Sars Cov 2 Pos Patient Lung Tissue Up
The Nlrp3 Inflammasome
Response To Elevated Platelet Cytosolic Ca2
Platelet Activation Signaling And Aggregation
Intestinal Immune Network For Iga Production
Inflammasomes
Graft Versus Host Disease
Normalized Enrichment Score
Xie St Hsc S1pr3 Oe Dn
Wu Silenced By Methylation In Bladder Cancer
Wp Zinc Homeostasis
Wp Cytoplasmic Ribosomal Proteins
Wong Embryonic Stem Cell Core
Wilensky Response To Darapladib
Weng Por Targets Global Up
Valk Aml Cluster 7
Tonks Targets Of Runx1 Runx1t1 Fusion Sustained In Granulocyte Up
Tonks Targets Of Runx1 Runx1t1 Fusion Granulocyte Up
Soga Colorectal Cancer Myc Up
Schuhmacher Myc Targets Up
Rashi Response To Ionizing Radiation 1
Pid Il8 Cxcr2 Pathway
Mootha Voxphos
Manalo Hypoxia Dn
Iritani Mad1 Targets Dn
Ichiba Graft Versus Host Disease 35d Up
Horton Srebf Targets
Eppert Ce Hsc Lsc
Casorelli Apl Secondary Vs De Novo Up
Type I Diabetes Mellitus
tRNA Aminoacylation
Translation
Srp Dependent Cotranslational Protein Targeting To Membrane
Sars Cov 2 Modulates Host Translation Machinery
Sars Cov 1 Modulates Host Translation Machinery
rRNA Modification In The Nucleus And Cytosol
RNA Processing
RNA Polymerase Ii Transcription Termination
Ribosome
Response Of Eif2ak4 Gcn2 To Amino Acid Deficiency
Respiratory Electron Transport Atp Synthesis By Chemiosmotic Coupling And Heat Production By Uncoupling Proteins
Respiratory Electron Transport
Nonsense Mediated Decay Nmd
Mitochondrial Translation
Mitochondrial Protein Import
Interferon Gamma Signaling
Eukaryotic Translation Initiation
Eukaryotic Translation Elongation
Cytosolic tRNA Aminoacylation
Citrate Cycle TCA Cycle
Cholesterol Biosynthesis
Asthma
Allograft Rejection
Activation Of The mRNA Upon Binding Of The Cap Binding Complex And Eifs And Subsequent Binding To 43s
Normalized Enrichment Score

### *shBCL11A + GC vs Cntrl shLuc + GC*

### *DAY 5*

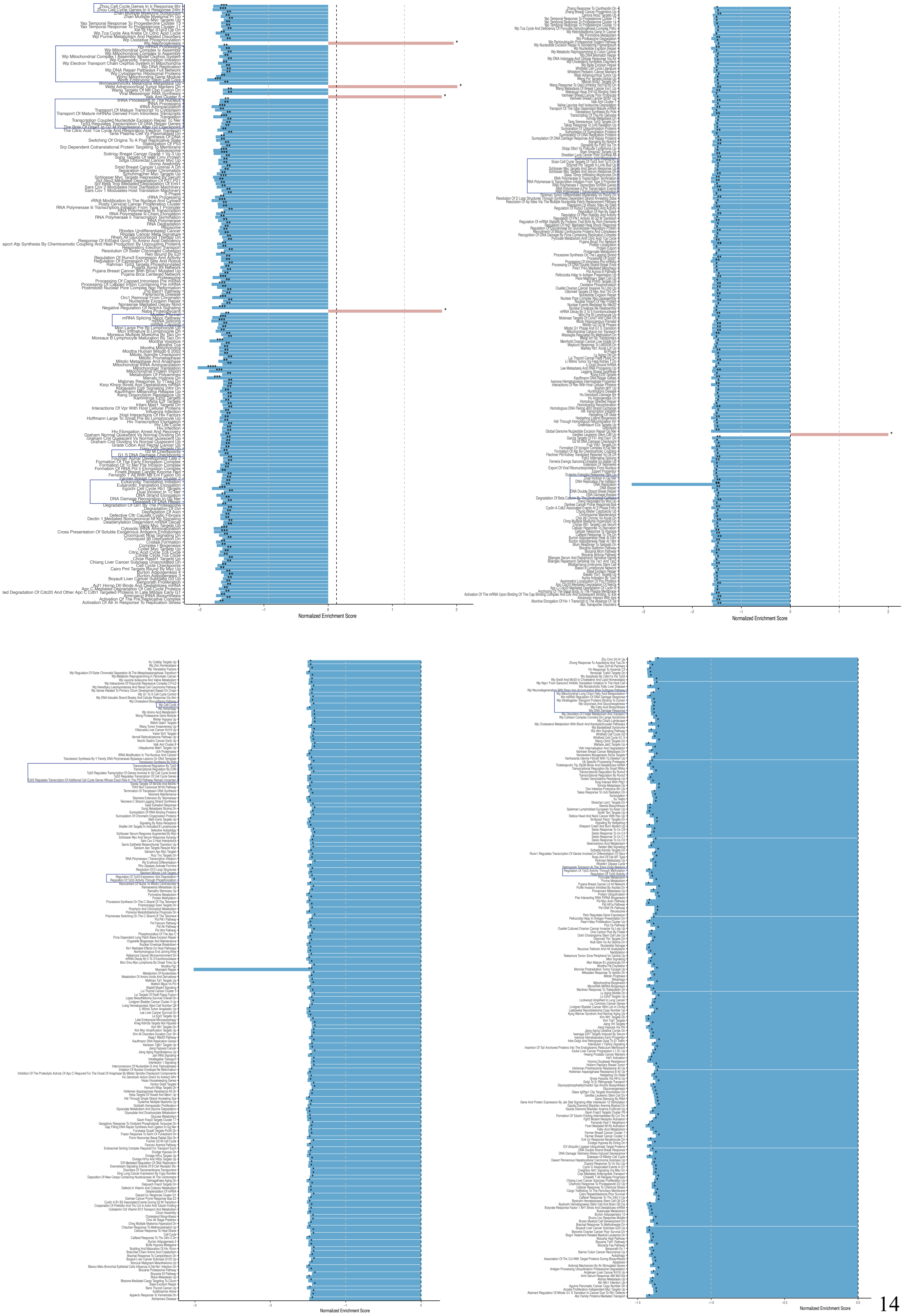

**Figure S7: GSEA enriched pathways analysis.** Comparison between control *shLuc* cells with or without GC at day 3 and day 5, between *shBCL11A* with or without GC at day 5, between *shBCL11A* and control without GC at day 5 and with GC addition at day 3 and 5. Red and blue columns indicate enriched and un-enriched pathway, respectively. All the data were compared using the HALLMARK gene set (h.all.v2023.1.Hs.symbols.gmt). Red and blue boxes indicate respectively up- and down-enriched pathways. FDR q-values among groups are indicated by asterisks (*, p<0.05; **,p<0.001; ***; p<0.0001****; p<0.00001).

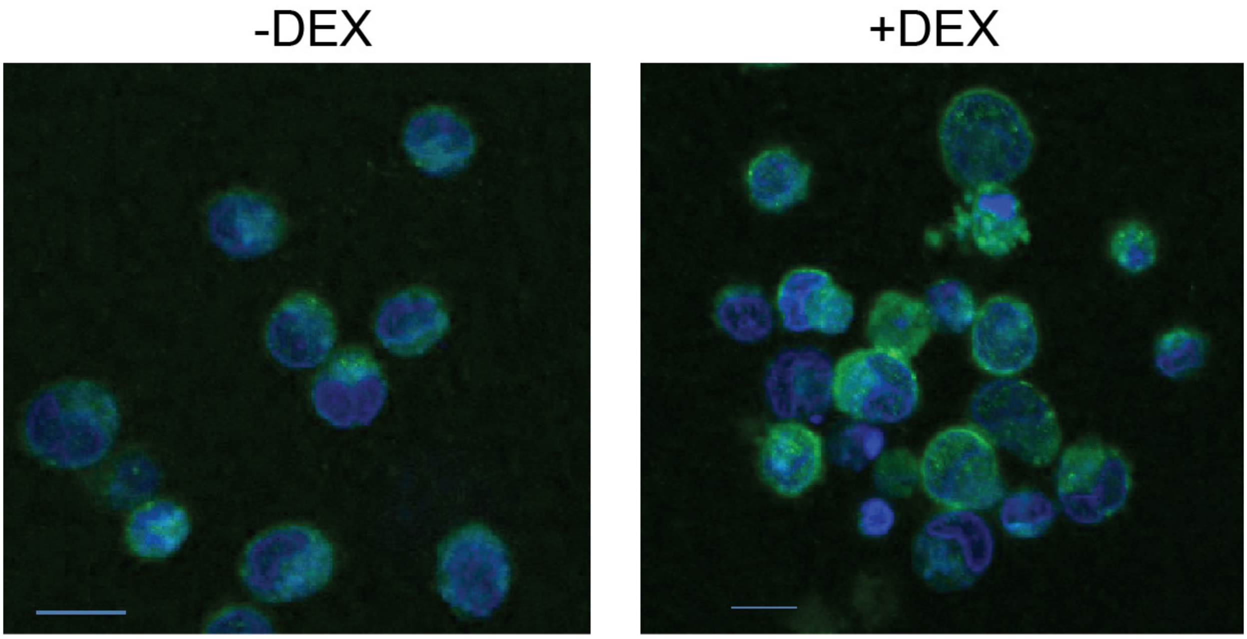


**Figure S8. Immunofluorescence analysis comparing the EKLF content in Erys generated by day 10 in cultures supplemented with or without Dex, as indicated.** Magnification 40X. A representative experiment is shown. The panel on the right is from [13]

**Table S1: Source of reagents, equipment and computer programs**

| | Catalog number | Provider |
|---|---|---|
| **Antibodies** | | |
| BCL11A | sc-33093 | Santa Cruz Biotechnology, Santa Cruz, CA |
| BCL11A | ab19487 | Abcam, Boston, MA |
| fluorescein isothiocya-nate-CD235a | clone JC159 | Agilent Dako, Santa Clara, CA, USA |
| GAPDH | CB1001 | Calbiochem, San Diego, CA |
| GATA1 | sc-1233 | Santa Cruz Biotechnology, Santa Cruz, CA |
| goat anti-mouse horseradish peroxi-dase-coupled second-ary antibody | 31430 | Invitrogen, Waltham, MA, USA |
| GRα | sc-8992H300 | Santa Cruz Biotechnology, Santa Cruz, CA |
| HBB | sc-21757 | Santa Cruz Biotechnology, Santa Cruz, CA |
| HBG | sc-21756 | Santa Cruz Biotechnology, Santa Cruz, CA |
| Histone 3 | ab10799 | Abcam, Boston, MA |
| Lamin B1 | Ab16048 | Abcam, Boston, MA |
| Pacific Blue 450-CD36 | 561535<br>(clone CB38) | BD horizon, Franklin Lakes, NJ, USA |
| **Chemicals** | | |
| Plasma from adult blood | H4522 | Sigma-Aldrich, Missouri, USA |
| CC110 supplements | 02697 | Stem Cell Techonologies, Inc., Vancouver, BC |
| Doxycyline | 72742 | Stem Cell Techonologies, Inc., Vancouver, BC |
| fetal bovine serum | SH30071.03IR25-40 | Hyclone, Logan, UT, USA |
| Ficoll-Hypaque | na | Amersham-Pharmacia Biotec, Uppsala, Sweden |
| heparin | 07980 | Stem Cell Techonologies, Inc., Vancouver, BC |
| Iscove's modified Dul-becco's medium | 30-2005 | Mascia Brunelli, Milan, Italy<br>ATCC, Manassas, VA,USA |
| L-Glutamine | 25030-081 | Gibco, ThermoFisher, Waltham, MA,USA |
| May-Grünwald-Giemsa | 63590 | Sigma-Aldrich, Missouri, USA |
| methylcellulose cul-tures | MethoCult | Stem Cell Techonologies, Inc., Vancouver, BC |
| methylcellulose cul-tures | MethoCult™ H4434 Classic | Stem Cell Techonologies, Inc., Vancouver, BC |
| Penicillin/streptomycin | SV30010 | Cytiva hyclone, Marlborough, MA, US |
| protamine sulfate | P4020-1G | Stem Cell Techonologies, Inc., Vancouver, BC |
| retronectin | T202 | Takara, Kusatsu, Shiga, Japan |
| StemSpan H3000 me-dia | 100-0073 | Stem Cell Techonologies, Inc., Vancouver, BC |
| StemSpan SFEM | 09650 | Stem Cell Techonologies, Inc., Vancouver, BC |
| Synperonic | 07579-250G-F | Sigma-Aldrich, Missouri, USA |
| transferrin | T0665-1G | Stem Cell Techonologies, Inc., Vancouver, BC |
| TRIzol | 15596026 | Invitrogen Life Technologies, Carlsbad, CA, USA |
| **Growth Factors** | | |
| dexamethasone | D1159-500MG | Sigma-Aldrich, Missouri, USA |
| Erythropoietin | 100-64-1MG | PeproTech, Cranbury, NJ, USA |

| | | |
|---|---|---|
| Estradiol | 1250008 | Sigma-Aldrich, Missouri, USA |
| G-CSF | 250-05 | Bouty SpA, Milan, Italy |
| Human recombinant insulin | SLCN8987 | Calbiochem, Darmstadt, Germany<br>Stem Cell Techonologies, Inc., Vancouver, BC |
| Hydrocortisone | 74142 | Stem Cell Techonologies, Inc., Vancouver, BC |
| Interleukin-3 | 200-03 | Biosource, San Jose, CA, USA<br>PeproTech, Cranbury, NJ, USA |
| Stem Cell Factor | 300-07-1mG | PeproTech, Cranbury, NJ, USA |
| **Molecular Biology Kits** | | |
| iScript Reverse Transcription Supermix | 1725282 | BIO-RAD, CA, USA |
| kit to amplify the three isoforms and the total *BCL11A* mRNA | Hs00250581_s1; Hs01093199_m1; Hs01093198_m1 Hs00256254_m1 | Applied Biosystems, Foster City, CA, USA |
| NE-PER nuclear and cytosolic extraction kit | 78835 | Thermo Scientific, Rockford, IL |
| QIAquick PCR purification kit | 28104 | Qiagen Sciences, Germantown, Maryland, USA |
| RNase OUT kit | 10777019 | Invitrogen Life Technologies, Carlsbad, CA, USA |
| RNeasy Micro Kit | 74004 | Qiagen, Hilden, DE |
| Trizol | 15596026 | Invitrogen Life Technologies, Carlsbad, CA, USA |
| **Retroviruses** | | |
| *BCL11A* shRNA vector (MISSION®shRNA) | clone # TRCN0000033449 | Sigma-Aldrich, Missouri, USA |
| shRNA control | | Made in the lab (see figure S2B for detail) |
| **Equipment** | | |
| 7700 Sequence 14 Detection System | na | Applied Biosystems, Foster City, CA, USA |
| Biohazard grade hood | 302611000 | Labconco, Kansas City, MO, USA |
| Cell Centrifuge | EP-5810A462 | Eppendorf SE, Hamburg, Germany |
| Cell Culture incubator | 51030284 | Thermo Scientific, Langeselbold, Germany |
| Cytocentrifuge | A78310250 | Shandon, Astmoor, UK |
| FACS Beckman Coulter | 16216213 | Becton Coulter Life Science, Indianapolis, IN, USA |
| Illumina Novaseq 6000 | 20012850 | Illumina, San Diego, CA |
| Light microscope equipped with a Coolsnap video camera | Axioscope | Zeiss, Oberkochen, Germany |
| Optical microscope | 473849 | Wild Heerbrugg, Switzerland |
| **Softwares** | | |
| FlowJo software | v7.6.4/v10 | Ashland, OR, USA |
| GENCODE v25 | na | na |
| Graph Pad9 Prism 9.5.1 | na | GraphPad Software, LLC, Boston, MA |
| GSEA software (4.3.1) | na | Broad Institute, Inc., Massachusetts Institute of Technology and Regents of the University of California |

| | | |
|---|---|---|
| KEGG gene-set | c2.cp.kegg.v2023.1.Hs.symbols.gmt | Broad Institute, Inc., Massachusetts Institute of Technology and Regents of the University of California |
| MSigDB | https://www.broadinstitute.org/gsea/msigdb/collections.jsp | Broad Institute, Inc., Massachusetts Institute of Technology and Regents of the University of California |
| R package DESeq2 | na | Bioconductor, Baden-Wuerttemberg, Germany |
| REACTOME gene-set | c2.cp.reactome.v2023.1.Hs.symbols.gmt | Broad Institute, Inc., Massachusetts Institute of Technology and Regents of the University of California |
| STARsolo (v27.9a) | na | GitHub, San Francisco, CA |
| Hallmark gene set | h.all.v2023.1.Hs.symbols.gmt | Broad Institute, Inc., Massachusetts Institute of Technology and Regents of the University of California |

*na= not available

**Table S2: Genes significantly regulated by GC both in control shLuc and sh*BCL11A* cells by day 5. Genes were considered upregulated and downregulates with a log2Fold Change > 0.500 (red fonts) or < - 0.500, respectively.**

| | Common significant genes | | | |
|---|---|---|---|---|
| | Cntrl shLuc: plus GC vs minus GC | | shBCL11A: plus GC vs minus GC | |
| **SYMBOL** | **log2Fold Change** | **padj** | **log2Fold Change** | **padj** |
| TFRC | 0.199 | 0.014 | 0.263 | 0.027 |
| LXN | 1.278 | 0.000 | 0.610 | 0.000 |
| FKBP5 | 1.187 | 0.000 | 1.335 | 0.000 |
| CA2 | 0.398 | 0.005 | 0.646 | 0.000 |
| PON2 | 0.654 | 0.001 | 0.698 | 0.023 |
| ID2 | 0.945 | 0.000 | 0.516 | 0.001 |
| TPST2 | 1.237 | 0.000 | 0.648 | 0.000 |
| CA1 | 0.442 | 0.005 | 0.775 | 0.000 |
| KIT | 0.786 | 0.000 | 0.561 | 0.039 |
| ALAS2 | -0.29 | 0.002 | -0.334 | 0.024 |
| PER1 | 2.034 | 0.000 | 2.049 | 0.000 |

**Table S3: Expression levels of genes from**[14]**in patients with congenic dyserythropoietic anemia (CDA) and WT control samples from**[15]**.** In blue, significantly down-regulated genes in CDA vs WT samples. Statistically significant genes are indicated in bold. The arrows indicate genes implicated in the self-replication of progenitor cells induced by GR activation identify by[14,16,17].

| **From** [14] | **From** [15] | | | | | | |
|---|---|---|---|---|---|---|---|
| **GENE ID** | **CDA d11** | **CDA d15** | **CDA diff5** | **WT d11** | **WT d15** | **WT diff5** | **T test (CDA vs WT)** |
| PNMT | 12.689 | 7.589 | 2.467 | 49.147 | 46.942 | 0.349 | 0.204 |
| RUSC2 | 2.320 | 3.975 | 5.550 | 1.335 | 5.362 | 4.012 | 0.814 |
| CFAP69 | 0.000 | 0.000 | 0.000 | 1.041 | 0.290 | 2.268 | 0.106 |
| SFRP4 | 1.501 | 2.891 | 0.000 | 0.037 | 0.278 | 0.698 | 0.259 |
| ALS2CL | 0.000 | 0.000 | 0.000 | 0.037 | 0.290 | 1.570 | 0.254 |
| DOCK3 | 0.273 | 0.000 | 0.000 | 0.245 | 0.568 | 6.804 | 0.316 |
| CCDC183 | 0.000 | 1.084 | 0.617 | 0.196 | 0.429 | 1.047 | 0.982 |
| MANSC1 | 0.273 | 0.000 | 3.700 | 6.566 | 4.239 | 4.361 | 0.057 |
| NOS1 | 0.000 | 0.000 | 0.000 | 0.012 | 0.000 | 11.688 | 0.373 |
| PLEKHH3 | 2.729 | 3.614 | 0.000 | 32.560 | 22.481 | 5.408 | 0.087 |
| **FAM179B** | **2.047** | **3.614** | **2.775** | **13.169** | **6.762** | **9.246** | **0.023** |
| NR4A1 | 1.092 | 7.589 | 0.000 | 1.886 | 0.921 | 8.374 | 0.815 |
| ZFP36L2 | 78.455 | 91.431 | 97.435 | 208.848 | 208.534 | 23.900 | 0.402 |
| CDKN1C | 2.183 | 5.782 | 0.000 | 0.551 | 2.195 | 0.523 | 0.427 |
| CD69 | 45.708 | 16.985 | 1.233 | 18.448 | 18.028 | 0.872 | 0.568 |
| PER1 | 22.513 | 19.876 | 5.858 | 134.038 | 65.449 | 4.187 | 0.243 |
| ZBTB41 | 2.456 | 7.228 | 5.550 | 4.618 | 5.992 | 7.153 | 0.621 |
| **TRPC1** | **0.136** | **0.000** | **0.000** | **1.127** | **1.022** | **1.570** | **0.002** |
| ZNF469 | 0.546 | 0.000 | 0.000 | 0.086 | 0.151 | 1.047 | 0.531 |
| **PPARA** | **4.639** | **1.446** | **5.858** | **23.814** | **17.472** | **29.483** | **0.006** |
| ZNF202 | 2.729 | 5.782 | 1.850 | 5.831 | 4.378 | 4.536 | 0.317 |
| CD4 | 21.558 | 41.560 | 94.043 | 3.577 | 28.915 | 4.187 | 0.158 |
| FER1L4 | 0.000 | 0.000 | 0.000 | 0.208 | 2.119 | 4.361 | 0.137 |
| CRY1 | 3.411 | 4.698 | 0.925 | 17.383 | 7.304 | 4.012 | 0.191 |
| **ZNF433** | **1.637** | **0.000** | **0.925** | **2.879** | **2.447** | **2.268** | **0.030** |
| TRIM62 | 3.275 | 4.698 | 2.158 | 2.205 | 2.309 | 0.523 | 0.144 |
| HCN3 | 0.000 | 0.000 | 0.000 | 1.825 | 1.476 | 0.174 | 0.082 |
| MCF2L2 | 0.000 | 0.000 | 0.000 | 0.478 | 0.467 | 21.632 | 0.346 |
| CASKIN2 | 0.000 | 0.000 | 1.233 | 0.024 | 0.429 | 1.745 | 0.653 |
| CASZ1 | 4.366 | 0.000 | 1.542 | 2.425 | 2.510 | 3.838 | 0.520 |
| **BCL11A** | **1.774** | **0.361** | **0.617** | **9.530** | **8.263** | **19.016** | **0.029** |
